# Micropatterned substrates to promote and dissect reprogramming of human somatic cells

**DOI:** 10.1101/111369

**Authors:** Jared Carlson-Stevermer, Ty Harkness, Ryan Prestil, Stephanie Seymour, Gavin Knight, Randolph Ashton, Krishanu Saha

**Author notes:** These authors contributed equally to this manuscript (Jared Carlson-Stevermer, Ty Harkness).

## Abstract

Reprogramming of human somatic cells to induce pluripotent stem cells (iPSCs) generates valuable precursors for disease modeling and regenerative medicine. However, the reprogramming process can be inefficient and noisy, creating many partially reprogrammed cells in addition to fully reprogrammed iPSCs. To address these shortcomings, we developed a micropatterned substrate that allows for dynamic live-cell microscopy of thousands of cell subpopulations undergoing reprogramming. Micropatterning facilitated a change in shape, size and clustering of nuclei to promote somatic identity erasure. Increased proliferation, cell density and decreased intercellular YAP signaling accompanied these nuclear changes. A combination of eight nuclear characteristics could be used to track reprogramming progression and distinguish partially reprogrammed cells from those that were fully reprogrammed.

Micropatterned substrates constitute a new tool for facile iPSC production and can be used in high-throughput to probe and understand the subcellular changes that accompany human cell fate transitions.

## INTRODUCTION

The cellular microenvironment and its engineering has recently received increased recognition as an important driver of mammalian cell fate decisions (Bratt-Leal, 2009; Hendrix et al., 2007; Iskratsch et al., 2014). In particular, rational use of surface modification and biomaterials synthesis has allowed for the tight control of biophysical cues presented to cells. This in turn has led to advances in precise control of phenotypes, such as stem cell self-renewal and differentiation (Bauwens et al., 2008; Wheeler et al., 1999). The physical microenvironment is sensed by cells through factors, such as talin (Lee et al., 2007; Vogel and Sheetz, 2006) and focal adhesion kinase (Mitra et al., 2005), which initiate cytoskeletal signaling cascades that act to regulate the structure of the nuclear lamina (Toh et al., 2015). The nuclear lamina is a powerful regulator of chromatin organization and dynamics and has been shown to play an active role in genome-wide gene expression by physically repressing genes in close proximity within the nucleus (Bronshtein et al., 2015; Guelen et al., 2008; Kind et al., 2015; Peric-Hupkes et al., 2010; Toh et al., 2015). Accordingly, there is a bidirectional, dynamic relationship between a cell’s gene expression and its biophysical state as regulated by its cellular microenvironment.

While known to be a major player in stem cell biology, (Dingal and Discher, 2014), microenvironmental regulation has been less studied in the context of somatic cell reprogramming to an induced pluripotent stem cell (iPSC) state. However, the ability to physically manipulate mechanotransduction genes leaves it as a prime candidate for engineering to increase reprogramming rate and efficiency. Notably, high resolution imaging of reprogramming cells has identified that nuclear geometry is dramatically altered during reprogramming (Cordie et al., 2014; Mattout et al., 2011). This may be due to the expression of kinases that activate cytoskeletal remodeling processes which are critical for reprogramming (Sakurai et al., 2014). These biophysical changes have traditionally been studied in the context of mesenchymal-to-epithelial transition (MET), a process that occurs relatively early during reprogramming (Li et al., 2010a; Liao et al., 2011; Samavarchi-Tehrani et al., 2010), concurrent with epigenetic changes indicating a loss of somatic identity, known as erasure (Gingold et al., 2014; Koche et al., 2011; Liao et al., 2011). Engineering strategies to alter the size and shape of reprogramming cells via culture on 3D hydrogels (Caiazzo et al., 2016; Choi et al., 2016) or PDMS micro channels (Downing et al., 2013) promotes reprogramming by activating genes associated with MET.

Another key player in microenvironmental signaling is the Hippo pathway, and the transcriptional coactivators, yes-associated protein (YAP) and transcriptional coactivator with PDZ-binding motif (TAZ). YAP and TAZ interact with Rho GTPase and the actin cytoskeleton to detect mechanical stress such as those induced by matrix stiffness or geometry of the area available for cell attachment (Sorrentino et al., 2014). In particular, localization of both factors is carefully regulated as they have specific functions in both the cytoplasm as well as the nucleus. In the cytoplasm YAP/TAZ binds to SMADs (Varelas et al., 2008), a key transducer of the TGF-β pathway that is antagonistic to reprogramming (Samavarchi-Tehrani et al., 2010; Woltjen and Stanford, 2009). When localized in the nucleus YAP and TAZ act as coactivators of the TEA domain (TEAD) family of transcription factors that influences several downstream targets. Many of these downstream targets are involved in cell proliferation and epithelial-to-mesenchymal transition (EMT), processes implicated in the erasure phase of reprogramming. Nuclear localization of YAP has also been identified as a key component to the maintenance of pluripotency in human pluripotent stem cells, while its exclusion to the cytoplasm can lead to differentiation (Lian et al., 2010; Sun et al., 2014).

Despite this progress in uncovering microenvironmental signaling during mammalian cell fate transitions, there is a current lack of understanding of cell-microenvironment interactions during the intermediate stages of reprogramming, precisely when reprogramming cultures are becoming heterogeneous. Although a decade has passed since the seminal reprogramming experiments, induction of iPSCs from human somatic cells continues to proceed at variable rates and can be partial, resulting in heterogeneous cell cultures (Buganim et al., 2012; Hanna et al., 2009, 2010; Smith et al., 2010; Zunder et al., 2015). Purifying complex mixtures of cells via subcloning and multi-week characterization of isolated subpopulations have the potential to lead to significant cell culture artifacts (Cahan and Daley, 2013) and increased costs (Paull et al., 2015; Yaffe et al., 2016). Cellular heterogeneity within a culture also makes mechanistic studies challenging. The high proportion of cells that do not undergo successful reprogramming means that single-cell assays profile many undesired cells, thus limiting the throughput of many assays (Buganim et al., 2012; Hanna et al., 2010; Zunder et al., 2015). Overall, understanding and managing population heterogeneity arising from reprogramming could result in significant gains in creating robust cell therapies and disease models.

Our approach dissects cell cultures undergoing reprogramming processes into ~10^3^ subpopulations. Unlike single-cell analysis, subpopulation analysis preserves the set of biochemical and biophysical cues presented to a cell by its microenvironment. Much of the work in the field to identify, evaluate, and enrich for high quality iPSCs has relied on dissociated cultures to perform flow cytometry or single cell sequencing (e.g., RNA-seq, ChIP-seq, ATAC-seq) at multiple time points (Buganim et al., 2012; Hansson et al., 2012; Mikkelsen et al., 2008; Polo et al., 2012; Zunder et al., 2015). Such methods disrupt the microenvironment, including cell-cell junctions, and drastically perturb the cytoskeleton, thus resulting in significant changes in the biophysical properties of cells undergoing reprogramming. Here, we utilize the microcontact printed well plate (μCP Well Plate), to enable multiplexed and independent control of the biophysical environment (Harkness et al., 2015). This technology allows us to perform high-content imaging experiments on hundreds of reprogramming μFeatures in a single well. We observe biphasic YAP signaling dynamics and understand how its spatial regulation affects erasure of somatic identity. Further, we gathered large, multidimensional data sets that are capable of characterizing reprogramming cell transitions and forming models that inform how cells react to their microenvironment during epigenetic reprogramming.

## RESULTS

### Controlled adhesion of reprogramming cells

We utilized a recently described cell culture platform, the microcontact printed (μCP) Well Plate, to gain control over cell adhesion during reprogramming (Harkness et al., 2015). The μCP Well Plate is formed by creating hydrophilic polyethylene glycol (PEG) brushes that resist protein adsorption at defined locations on a gold coated glass sheet. This sheet is then combined with a standard tissue culture well plate to form a μCP Well Plate (Figure 1—figure supplement 1A). When seeded with Oct4-Sox2-Klf4-cMyc reprogrammable fibroblasts (Cordie et al., 2014) and upon induced reprogramming factor expression, this platform constrained cell-to-cell contacts and controlled the geometry of cellular aggregates (Figure 1A, left). We are able to pattern cells into various geometries and within microfeatures (μFeatures) of a defined diameter. In addition, μCP Well Plates allowed for high-content imaging and permitted a high degree of multiplexing on a single plate.

**Figure 1.**
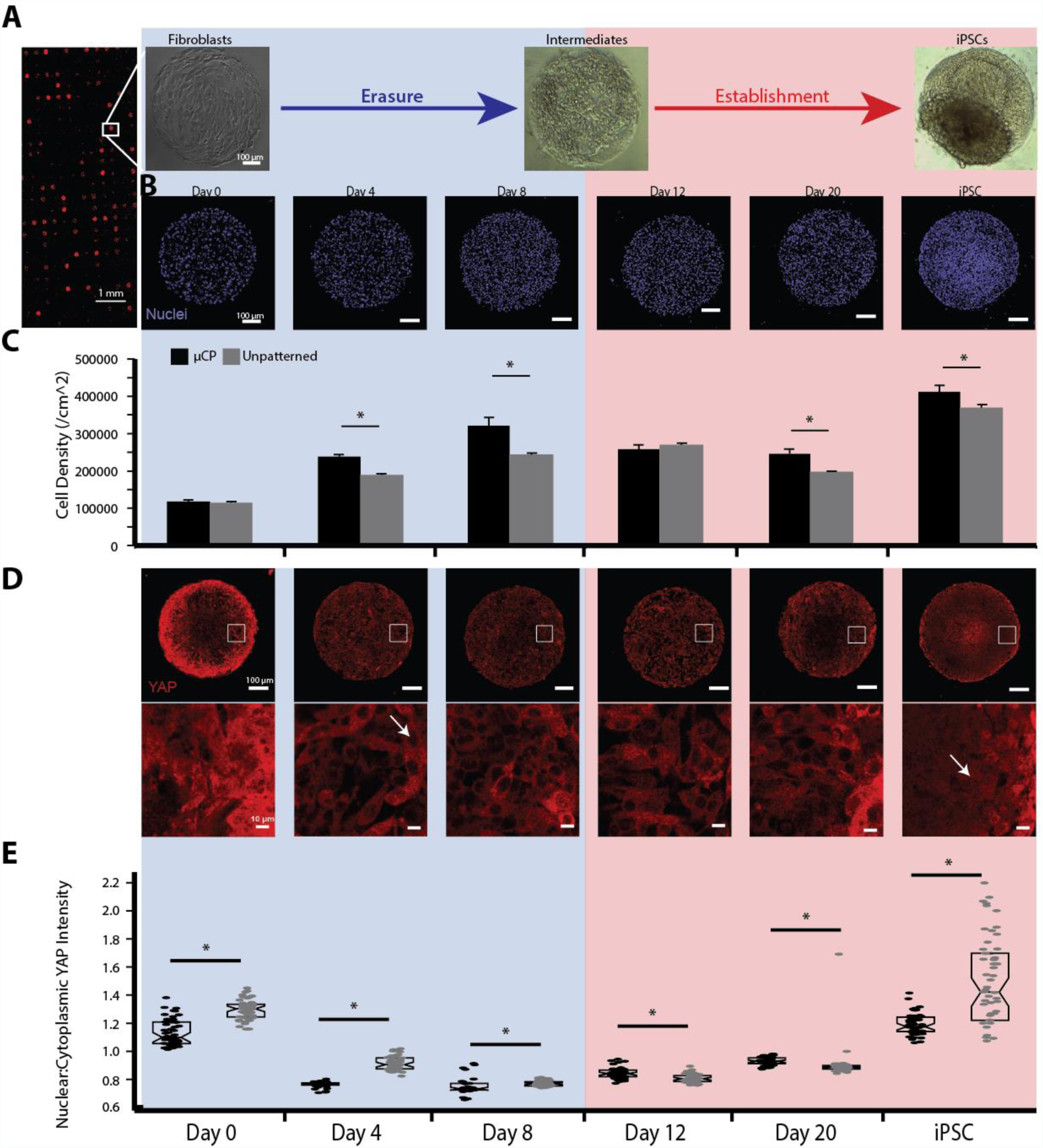
Reprogramming fibroblasts on micropatterned substrates causes erasure of somatic cell identity by increasing cell density and proliferation, and modulating YAP localization. **A)** Left: hPSCs expressing H2B-mCherry on the μCP Well Platform. Right: Model showing the erasure of somatic cell identity and the accompanying markers to an intermediate cell fate followed by the establishment of pluripotency. **B)** Representative images of the progression of a cell aggregate on a μCP Well Plate through a reprogramming time course. **C)** Cell density through reprogramming on micropatterned and unpatterned substrates. Micropatterned substrates had higher cell densities during erasure (D8 and D12) (n=49 technical replicates, mean ± 95% CI, Student’s two-tailed t-test, \**p*<0.05). **D)** YAP staining of reprogramming time course. In fibroblasts, YAP is evenly distributed through the cell. During erasure YAP is excluded from the nucleus and slowly moves back into the nucleus during establishment. Bottom: Magnified image of white box in middle row. **E)** Quantification of the ratio of YAP intensity in the nucleus to the cytoplasm on μFeatures vs unpatterned substrates. μFeatures had significantly less YAP in the nucleus throughout erasure (n=49 technical replicates, mean ± 95% CI, Student’s two-tailed t-test, \**p<0.05*).

We next assessed the ability of the μCP Well Plate to sustain long-term reprogramming studies. The high-content imaging capabilities enabled us to track large numbers of cell subpopulations (>100 μFeatures per well in a 24 well plate) at multiple time points with high reliability of observing the same subpopulation within each μFeature from time point to time point (Carlson-Stevermer et al., 2016). Reprogramming cells remained viable, attached, and confined to the desired micropatterns and the establishment of the pluripotency network over a 3-4 week time course (Figure 1B). In separate experiments that co-cultured pluripotent stem cells (PSCs) with fibroblasts, we ensured strong adhesion of each cell type for more than one month (Figure 1—figure supplement 1B), thus, one cell type does not displace or cause detachment of one cell type over another. Using a previously established protocol, we were also able to track live reprogramming μFeatures with antibodies targeting cell surface markers identifying fibroblasts and pluripotent cells (Chan, 2009; Cordie et al., 2014; Quintanilla et al., 2014) (Figure 1—figure supplement 1C) and found that fibroblasts (CD44^+^/TRA-1-60^-^), iPSCs (CD44^-^/TRA-1-60^+^), and intermediate (CD44^-^/TRA-1-60^-^) cells were readily detected on the same μFeature. These changes correspond to two distinct phases of reprogramming, the erasure of somatic cell identity to an intermediate cell state followed by the establishment of pluripotency (Figure 1—figure supplement 1C). These results demonstrate that the micropatterned substrate could impose strong physical constraints on each population over the entire course of reprogramming.

### Biphasic cell density and YAP activity

Because we observed dramatic changes in cell clustering by the end of our reprogramming experiments, we performed immunocytochemistry at intermediate time points to monitor cell organization. Cell density on both micropatterns and standard substrates increased during erasure, peak at the end of erasure, and then decreased during establishment (Figure 1C). The peak cell density on micropatterns occurred more quickly than on standard substrates (day 8 vs. day 12). Further, PSCs on micropatterns were denser than on standard substrates. Consistent with higher cell densities during erasure, the percentage of proliferating cells on the μCP Well Plates was higher than on standard, unpatterned substrates (Figure 2—figure supplement 1A).

We also profiled a key component of the Hippo pathway, YAP, as this pathway plays a role in mechanotransduction and is an important sensor of cell density (Aragona, 2013; Dupont et al., 2011; Li et al., 2010b; Low et al., 2014). Additionally, it is crucial for stem cell maintenance (Dupont et al., 2011; Hsiao et al., 2016; Lian et al., 2010; Musah et al., 2014), and is a regulator of epithelial-to-mesenchymal transitions (EMT) (Liu-Chittenden et al., 2012; Xie et al., 2013; Zhao et al., 2008), which has been implicated in the erasure phase of reprogramming. Nuclear YAP localization, a hallmark of active Hippo signaling, was biphasic, where minimal nuclear levels are seen more quickly on micropatterns than on standard substrates (day 4 and 8 vs. day 8 and 12) (Figure 1D). Within a μFeature for all cell types, YAP levels were higher at the perimeter of the μFeature (Figure 2—figure supplement 1B), but YAP levels averaged across each entire μFeature were biphasic. In fibroblasts, YAP was expressed at high levels and was evenly spread throughout the nucleus and cytoplasm. While initial cell densities were equivalent between micropatterned and standard substrates, YAP nuclear localization was already decreased on day 0 on the micropatterns. Low nuclear YAP levels were seen during erasure and YAP began to translocate back into the nucleus during establishment (Figure 1E). In PSCs, YAP translocation back to the nucleus can sustain pluripotency at lower levels than what is needed on standard substrates (1.2 vs. 1.5 in Figure 1E). During erasure, higher cell density on micropatterns led to low cytoskeletal tension and decreased YAP activity. This decrease in YAP activity, however, did not persist during establishment. For the final establishment state, micropatterning lowers the bar for YAP activity leading to successful reprogramming.

### Hyperactive TAZ slows erasure

To understand the effects of the Hippo pathway on reprogramming on our micropatterns, we infected our reprogrammable fibroblasts with a hyperactive TAZ (hTAZ) lentivirus prior to seeding on the μCP Well Plate platform. hTAZ (Yang et al., 2014) contains a serine residue (S89) that has been mutated to alanine, which prevents phosphorylation at this residue. Without phosphorylation, TAZ remains in the nucleus and continues to transcribe TEAD genes. We observed that cell density decreased significantly during the erasure phase of hTAZ populations and took an additional 16 days for the cell density to reach the 1000 cells/feature density observed in the control at day 8 (Figure 2A). We found that YAP remained in the nucleus at a higher level than control cells during the first 20 days of factor expression (Figure 2B) before decreasing to control levels after 24 days. Concurrently, the mean amount of YAP within cells significantly increased before returning to day 8 control levels by day 24, supporting that hTAZ upregulates the Hippo pathway during the erasure phase (Figure 2C). We also measured the mean intensity of Snail, a marker of EMT — a transition that is opposite of the mesenchymal-to-epithelial (MET) transition previously described during the erasure phase. Snail levels were increased in hTAZ samples at day 8 (Figure 2D, Figure 2—figure supplement 1C). With additional culture to day 20 and day 24, Snail levels eventually decreased (Figure 2—figure supplement 1D). Further, transduced cells had elevated levels of CD44 at day 8 (Figure 2E), indicating that these cells had retained - and not erased somatic identity - to the same extent as in the control cells. We did not observe any Nanog+ cells during the 24 days of reprogramming with the hTAZ cells. Taken together, hyperactivity of the Hippo pathway counteracts the effects of micropatterning during erasure.

**Figure 2.**
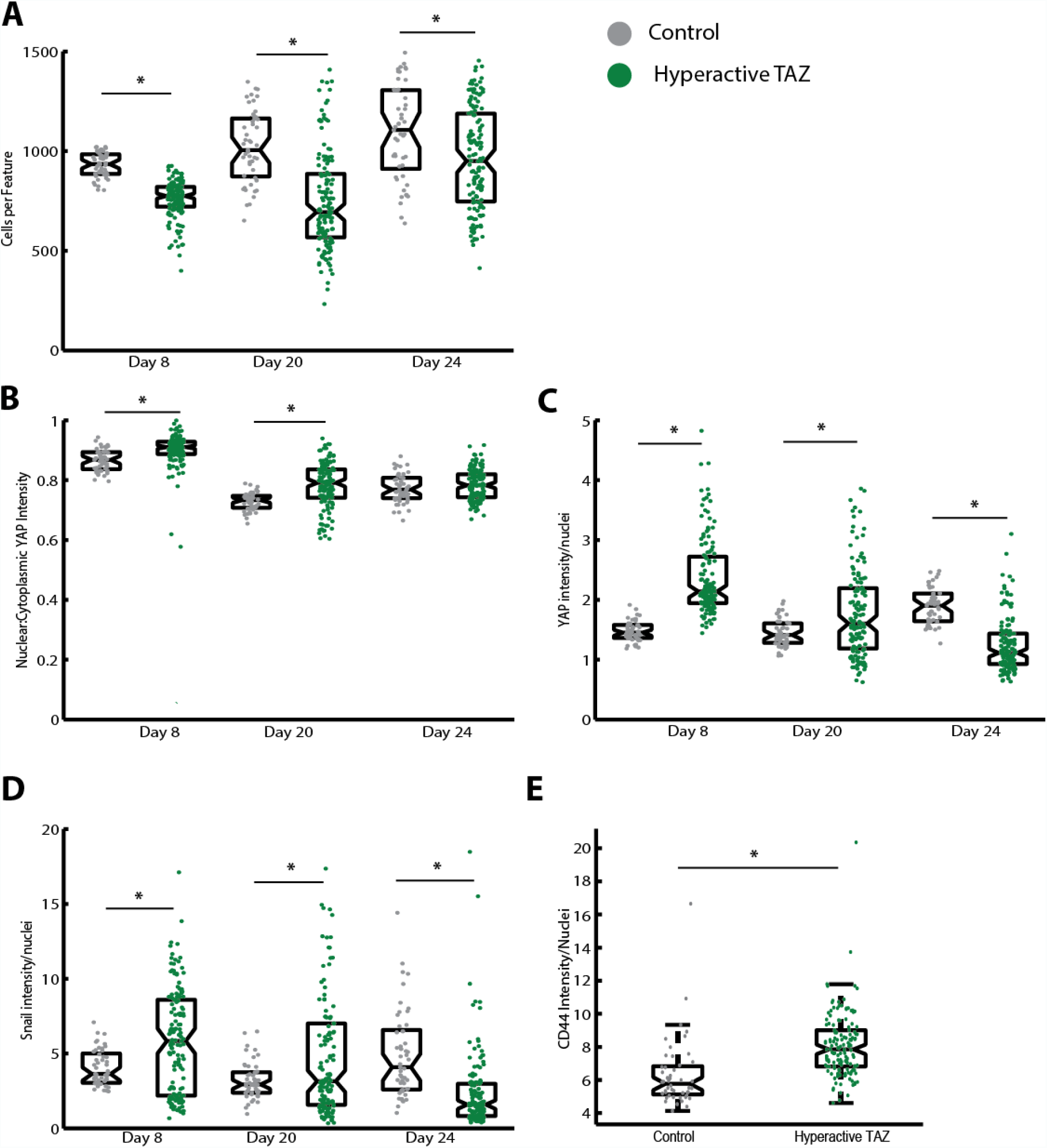
Hyperactive TAZ slows the erasure of somatic cell identity. **A)** Control μFeatures have increased cell density compared to hyperactive TAZ μFeatures. Proliferation and cell number are early indicators of reprogramming. **B)** Quantification of nuclear to cytoplasmic YAP intensity on μFeatures. Hyperactive TAZ causes increased nuclear localization during the first 20 days of reprogramming. **C)** Hyperactive TAZ upregulates YAP expression in reprogramming cells until day 24 when it significantly decreased. **D)** Snail protein is regulated similarly to YAP when expressing hyperactive TAZ. Control μFeatures express Snail at a constant level. **E)** CD44 expression after 8 days in cells following standard reprogramming (grey) or infected with hyperactive TAZ lentivirus (green). Hyperactive TAZ slows erasure of somatic cell identity (n=49 technical replicates, mean ± 95% CI, Student’s two-tailed t-test, \**p<0.05*).

### Establishment on micropatterned substrates

We next wanted to explore the effect of our μCP Well Plates on the establishment of pluripotency. When μFeatures were stained for Nanog expression, a marker of fully reprogrammed iPSCs, we found that there was a wide range of expression levels within a single μFeature (Figure 3A). Furthermore, due to the 3D nature of reprogrammed μFeatures we confirmed that expression levels were consistent within a μFeature at all distances away from the substrate (Figure 3—figure supplement 1A, B). Interestingly we found that there was a significant relationship between the area of the circular μFeature and the efficiency at which the entire μFeature reprogrammed. When the μFeature diameter was greater than 450 μm, the clear majority of μFeatures expressing Nanog expressed it in only less than half of the cells (Figure 3B). However, when the μFeature diameter was less than 450 μm, all of the μFeatures that expressed Nanog expressed it in over half of all cells (Figure 3B). This suggests there may be some inherent characteristic of area available for cell attachment that influences efficiency. It has previously been shown that confinement of cells into different geometries causes different stress patterns and subsequently differing downstream effects (Jain et al., 2013). To explore this effect, we modulated the geometry of μFeatures on our plates (Figure 3—figure supplement 1C) and measured the number of μFeatures expressing Nanog at any level. We found that there was no statistically significant difference between any of the geometric shapes (Figure 3C). We next hypothesized that the number of cells seeded initially may influence reprogramming efficiency. We found that this was the case up to a point where increasing cell number no longer had any effect on the number of μFeatures expressing Nanog (Figure 3D). This may be caused by the fact that fibroblasts can only cluster so tightly before they no longer attach to the confined area of the μFeatures. Increased cell number up to this point may help with clustering and erasure of somatic cell identity.

**Figure 3.**
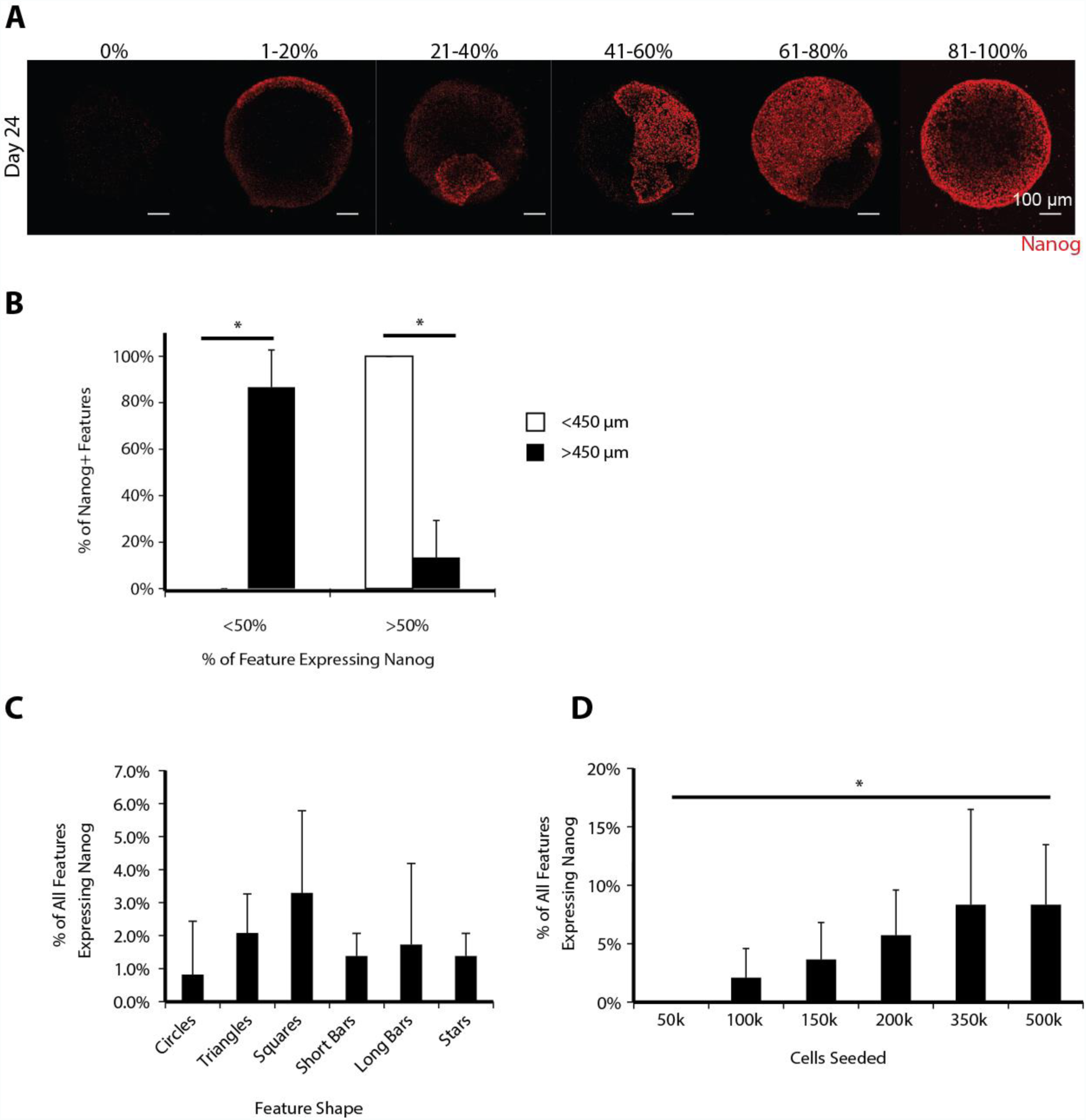
Reprogramming efficiency on μCP Well Plates is affected by cell density and μFeature size but not by geometry. **A)** Representative images of μFeatures classified into 20% Nanog+ bins. **B)** Reprogramming efficiency by size of μFeature. When μFeature size was greater than 450 μm most μFeatures had less than 50% of cells express Nanog at the end point. In comparison, when μFeature size was than 450 um, 100% of μFeatures had over half of the μFeature expressing Nanog (n=3 biological replicates, mean ± 95% CI, Student’s two-tailed t-test, *p*=0.001, 0.002). **C)** Percent of μFeatures expressing any level of Nanog on six different shapes. There was no significant difference in reprogramming efficiency between shapes (n= 3 biological replicates, 27 technical replicates each, mean ± 95% CI, One-way ANOVA, *p*=0.46). **D)** Percent of μFeatures expressing Nanog as a function of seeded cell density. Reprogramming efficiency increased up to a maximum where there was no longer room for additional cell attachment (n=4 biological replicates, 24 technical replicates each, mean ± 95% CI, One-way ANOVA, *p*=0.037).

### Nuclear characteristics to track reprogramming

In our immunocytochemistry images, we noted dramatic changes in shape and size of nuclei throughout the reprogramming process. We hypothesized that these changes could be used as a tool to track progression through the two phases of reprogramming. As a proof of concept, we seeded three distinct cell types on to the μCP Well Plates: PSCs, fibroblasts, and neural progenitor cells (NPCs) and used a Hoechst dye to track nuclear characteristics through high-content imaging techniques. These images were then fed through a CellProfiler (Carpenter et al., 2006) pipeline to identify nuclei within the images and output a set of geometrical, intensity, and clustering measurements (Figure 4—figure supplement 1A). We started with a large dataset of 32 nuclear characteristics that was then filtered to a set of 8 core characteristics by evaluating correlation between measured variables. These core characteristics are: area, perimeter, mean radius, nuclear shape index (NSI), extent, solidity, nearest neighbor, and number of neighbors (Figure 4—figure supplement 1B). We initially attempted to identify cell type by analyzing individual cells using two methods: principal components analysis (PCA) and t-distributed stochastic neighbor embedding (t-SNE) (Maaten and Hinton, 2008). However, neither method could faithfully distinguish the three different cell types from each other (Figure 4—figure supplement 1C). Based off results from tracking YAP dynamics, we next turned to analyzing nuclear characteristics on a per-μFeature basis using the same methods. Cell types analyzed on an μFeature level separated cleanly using both methods but were more highly clustered using PCA (Figure 4—figure supplement 1D). We proceeded with this method for future models.

To test our hypothesis that reprogramming cells can be tracked using nuclear characteristics, we stained reprogramming cells, at four intermediate time points (day 4, 8, 12, and 20) as well as at the beginning and end of the experiment. When core characteristics for each time point were used to form a PCA model, a progression from fibroblasts to iPSCs emerged when using three principal components (PC) (Figure 4A). We found that there was a biphasic progression in the first two principal components while the third principal component (PC3) progressed in a linear fashion (Figure 3B). Combining all three PCs explained 95% of the variance in the model (Figure 4—figure supplement 2A) and resulted in the formation of a spiral tracing the progression of cell state from fibroblast through erasure to intermediate and finally establishment to iPSC (Figure 4A). By analyzing the loadings of each PC we found that PC1 is largely driven by variables describing nuclear shape (extent, perimeter, solidity, NSI) while PC2 corresponds largely to changes in nuclear size (area, mean radius) (Figure 4— figure supplement 2B). In contrast, PC3 is dominated by shifts in nuclear clustering (nearest neighbor, number of neighbors). This result is supported by visual inspection where fibroblast nuclei are elongated and far apart from each other whereas PSC nuclei are circular and close together (Figure 1B).

**Figure 4.**
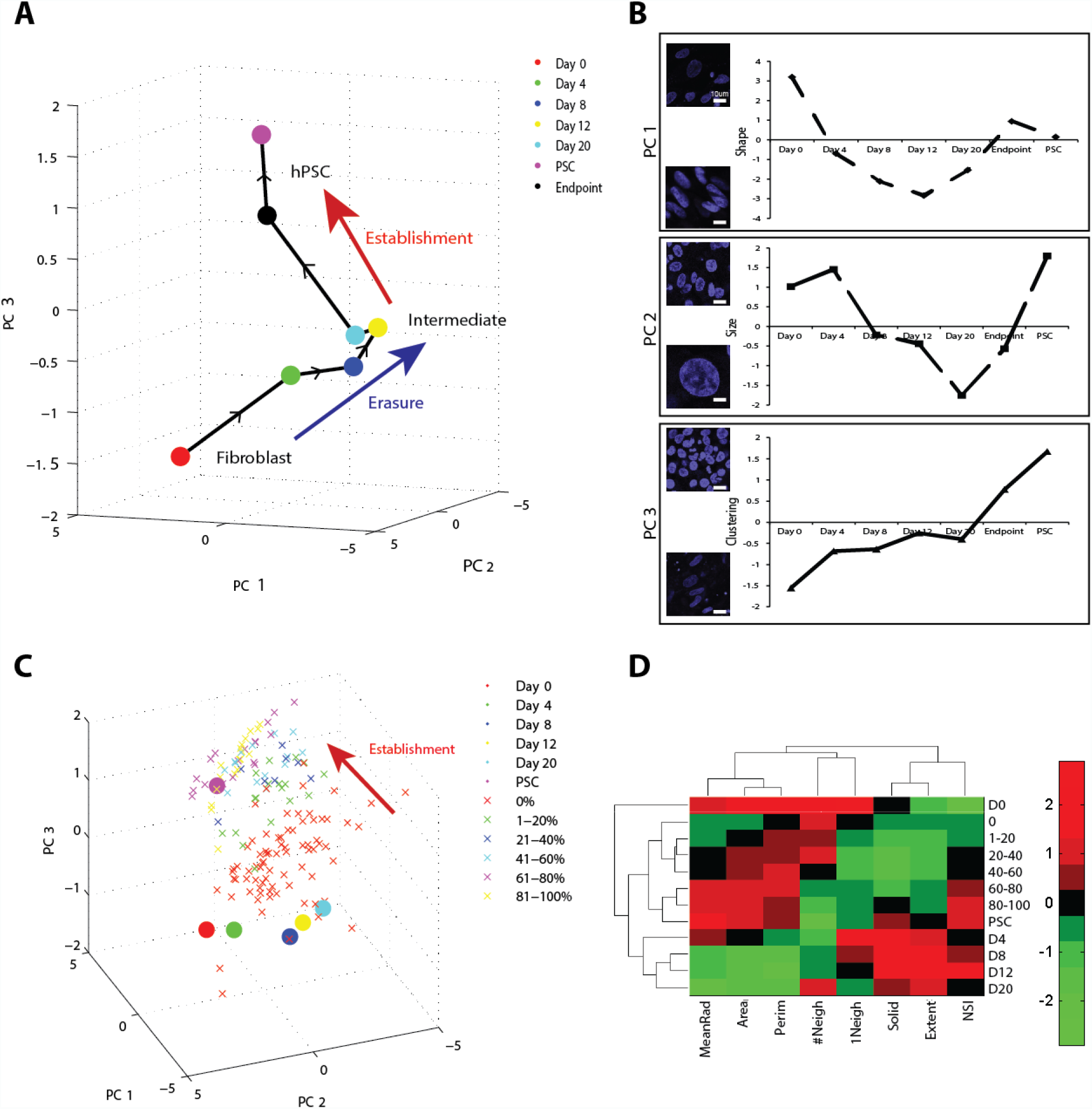
Separation of reprogramming intermediates via nuclear characteristics. **A)** Principal components analysis of subpopulation nuclear measurements on human fibroblasts undergoing microscale reprogramming. Centroid values for each reprogramming time point indicate a 3D spiral of reprogramming progression corresponding to erasure and establishment phases of reprogramming are shown. Model was generated using 49 replicates at each time point. **B)** Breakdown of centroid values across all three principle components. **C)** PCA model as in (A). Endpoint data is color coded by %Nanog expression. Low Nanog^+^ μFeatures were similar to intermediate time points while high Nanog^+^ μFeatures clustered close to PSCs. **D)** Hierarchical clustering of each time point in reprogramming and endpoint data classified into 20% bins of Nanog+ μFeatures. 60-100% positive μFeatures clustered closely to PSCs while reprogramming intermediates shared nuclear characteristics. Seeded fibroblasts prior to reprogramming (D0) were unlike any other cell type. Clustering based on nuclear characteristics is similar to what was found via PCA loadings.

The largest shifts in PC space occur between days 0-8 corresponding with erasure of somatic cell identity. Temporally, this shift matches closely with the well-studied MET that occurs within the first 10 days of reprogramming and involves dramatic shifts in cell morphology. Following erasure there was a clustering of time points between Day 8-20 that corresponds to the rise of an intermediate cell type (CD44^-^/TRA-1-60^-^) (Figure 1—figure supplement 1C). A final transition corresponding to establishment of pluripotency occurred between day 20 (light blue) and the endpoint (black) of the experiment. Nuclei following establishment are highly circular, densely packed and cluster closely to hPSCs (purple) in PC space. We noticed that there was significant heterogeneity in the endpoint populations with different μFeatures clustering closely to various time points in the reprogramming process. To confirm this heterogeneity we fixed and stained endpoint μFeatures for Nanog and found that there was a wide range of successfully reprogrammed μFeatures (Figure 3A). When endpoint μFeature data was plotted in PC space μFeatures that showed no reprogramming (0% Nanog^+^) clustered closely to intermediate cell states. Conversely, μFeatures that completely reprogrammed (81-100% Nanog^+^) clustered with hPSCs (Figure 4C).

To validate our PCA map, we used hierarchical clustering to determine the distance between both experimental time points as well as variables. As seen in the PCA map, solid, extent, and NSI are closely related as are number of neighbors and closest neighbor (Figure 4D). Interestingly, area and perimeter are the closest related variables after hierarchical clustering but logically are directly proportional when describing the physical properties. We next validated the reprogramming spiral generated from the PCA map. As seen in the map, day 0 fibroblasts are the farthest away from any other population. Tracing up the spiral, day 8, 12, and 20 cluster the closest together of the reprogramming intermediates followed by day 4. Endpoint μFeatures clustered closer to hPSCs than any of the reprogramming intermediates. This suggests that many, if not all μFeatures may have started the establishment of pluripotency but had a longer latency period in the intermediate cell stage. Of the endpoint data, μFeatures that were 60-100% Nanog^+^ clustered closely to hPSCs. Conversely, μFeatures that were 40-60% Nanog^+^ were more similar to μFeatures that did not contain any Nanog than hPSCs.

We further transformed the nuclear measurements from the hTAZ experiments into PC space defining the reprogramming spiral (Figure 4—figure supplement 2C). We found that day 8 of the hTAZ and day 20 data were most similar to day 4 reprogramming intermediates. This supports the conclusion that hTAZ slowed erasure. Additionally, when day 24 data was analyzed it appeared most similar to day 20 μFeatures. This suggests that while cells overexpressing TAZ protein can undergo erasure, the latency period is increased and cells may never undergo establishment. We next performed hierarchical clustering on reprogramming time points including hTAZ μFeatures. The reprogramming clustering diagram slightly changed with the addition of this data. Clustering of cell populations remained the same as in the previous model while hTAZ μFeatures clustered separately (Figure 4—figure supplement 2D).

### Predicting reprogramming using nuclear characteristics

If analysis of nuclear characteristics can separate reprogramming intermediates from one another, we reasoned that these same measurements would be able to predict which μFeatures were successfully reprogrammed at the end of the time course. Creation of this model could be used to aid in the rapid purification of high purity iPSCs. We created a partial least squares regression model (PLSR) using the eight minimal nuclear characteristics described previously and output the expected fraction of each μFeature expressing Nanog. The resulting model explained 78% (Figure 5A) of the variance in endpoint reprogramming culture and relied heavily on NSI and nearest neighbor characteristics (Figure 5B), hallmarks of pluripotent cells. The model was highly predictive of reprogramming efficiency with a root mean square error of prediction (RMSE) of 0.13 (Figure 5C). We compared this model to a second PLSR model using immunostaining of TRA-1-60 levels as the experimental data. This second model was not as predictive of complete reprogramming, possessing a RMSE of 0.28 (Figure 5D). We confirmed that TRA-1-60 expression was not wholly predictive of Nanog expression by staining for both markers and observed significant difference between the two populations (Figure 5E).

**Figure 5.**
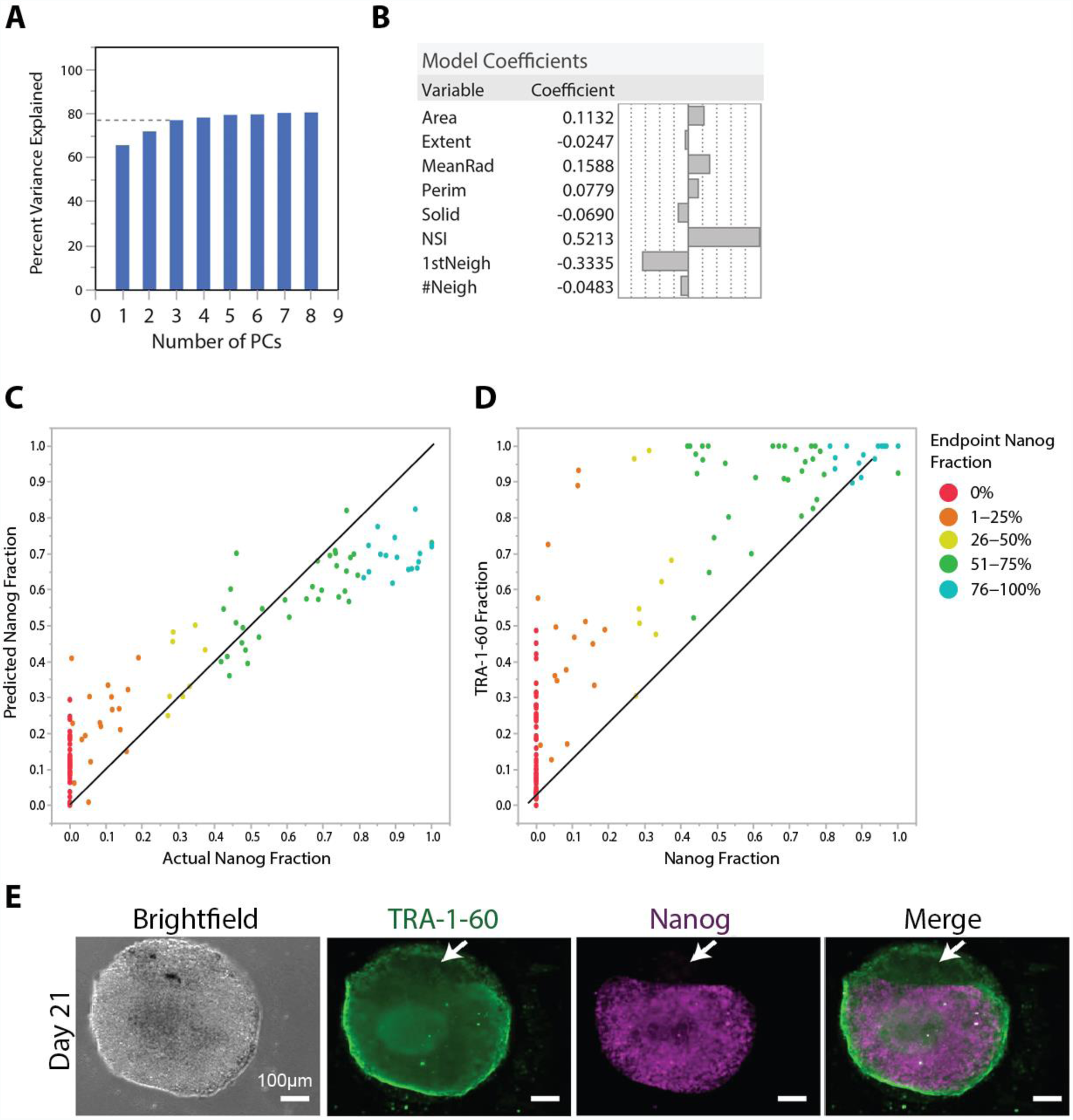
Nuclear characteristics are predictive of reprogramming. **A)** Percent variance explained with increasing number of principle components in PLSR model. 3 principle components explained over 75% of the data while additional PCs showed limited returns. **B)** Model coefficients of PLSR model. Nuclear shape and nearest neighbor heavily influenced the model **C)** PLSR model predicting Nanog^+^ fraction of μFeatures using only nuclear characteristics. Data points are individual μFeatures color coded by %Nanog positive. Model was predictive of reprogramming with low error (RMSE=0.13). **D)** PLSR model predicting Nanog^+^ fraction using TRA-1-60 staining as input variable. Model was less predictive (RMSE=0.28) than use of nuclear characteristics. **E)** Representative image showing difference between TRA-1-60 expression and Nanog expression.

### Isolation of high quality iPSCs

The terminal goal of any reprogramming platform is to successfully isolate iPSCs that can be used for downstream applications. Overall, reprogramming on μCP Well Plates enabled the simple derivation of pure iPSC lines with minimal time and effort spent on purification. The physical separation of micropatterns from one another, combined with high fraction of Nanog expressing cells, even up to 100% throughout the μFeature (Figure 6A) resulted in easy picking and isolation of reprogrammed cells. However, we observed that even the picking of impure colonies based on early TRA-1-60 expression (Figure 6B) resulted in the presence of Nanog^+^ cells that could be isolated following one additional round of picking (Figure 6C-D). We confirmed that these cells expressed Nanog at the same level as ESCs using flow cytometry (Figure 6F). To stringently assess the pluripotency of established lines we cultured cells isolated from 2 separate μFeatures for 10+ passages in fully-defined stem cell media and then formed embryoid bodies (EBs) using the AggreWell method. iPSCs and EBs were then subjected to the TaqMan**™** ScoreCard assay, a benchmarked quantitative assay for pluripotency (Tsankov et al., 2015). iPSCs had a similar expression profile to nine control hESC lines and EBs expressed genes indicative of all three germ layers (Figure 6G, Figure 6—figure supplement 1). These results indicate that reprogrammed cell lines isolated from the μCP Well Plate are fully pluripotent.

**Figure 6.**
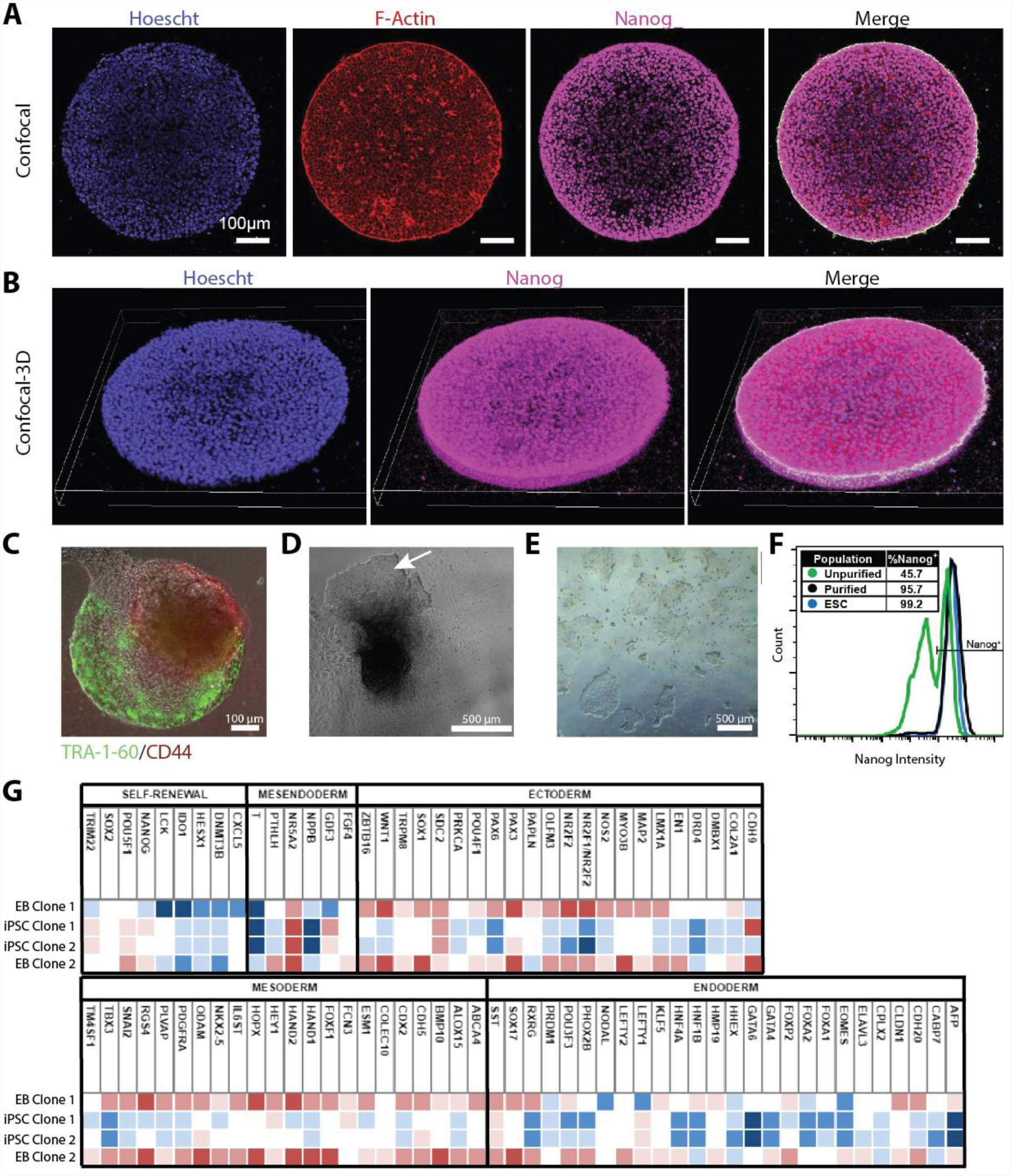
μCP Well Plates enable isolation of pure iPSC populations. **A)** Confocal image of μFeature that underwent complete reprogramming. **B)** 3-D image stack showing reprogramming occurred across entire μFeature **C-E)** Representative images of the derivation of pure iPSC lines from μFeatures. **C)** Day 21 live cell μFeature live-stained with CD44 (red) and TRA-1-60 (green) antibodies. **D)** Aggregate from (C) three days after picking from a μFeature. White arrow represents expanding iPSC colony. **E)** DXC-independent iPSC line after one picking step from (D) and several passages in standard pluripotent culture conditions. **F)** Flow cytometry histogram indicating the percentage of Nanog+ cells before (D) and after (E) one purification step. **G)** Taqman Scorecard^TM^ of iPSCs and differentiated embryoid bodies (EB) from microscale reprogramming. Two iPSC clones expressed pluripotency factors at the same level as nine benchmarked PSC lines. EBs generated from iPSC lines expressed high levels of genes for all three germ layers, significantly above the benchmarked PSC lines. Blue boxes are downregulated while red boxes are upregulated compared to 9 control hPSC lines.

## DISCUSSION

Here we present a micropatterned substrate for the generation of human iPSCs. The starting fibroblasts, intermediate cells, and endpoint fully reprogrammed cells all adhere onto these substrates with minimal detachment, thereby enabling novel control over the microenvironment during all stages of reprogramming of human fibroblasts. For human whole blood reprogramming, these substrates also permit adhesion of intermediate and endpoint iPSCs (data not shown), which could be used to control the establishment stage of reprogramming. Because skin and blood constitute the majority of cell sources for research and clinical-grade reprogramming projects and biobanks (Kreiner and Irion, 2013), these substrates have strong potential to advance the manufacturing of iPSCs for a variety of disease modeling and regenerative medicine applications.

The micropatterned substrate importantly enables the non-destructive optical tracking of subcellular changes with these populations without disrupting cell-cell contact or the microenvironment. Thus, longitudinal study of single cells and structures within them produce large datasets that can be analyzed for mapping cell fate transitions and trajectories with reprogramming cultures. The maps generated by our approach reflect the complexity seen by flow and mass cytometry in mouse reprogramming cultures (Mikkelsen et al., 2008; Zunder et al., 2015). While mouse and human reprogramming have important differences, nearly all studies describe erasure and establishment or maturation of pluripotent stem cells. A large variety of intermediate states were also identified by flow and mass cytometry, which is consistent with the heterogeneity seen in our maps, notably after erasure and during establishment. Noise in reprogramming during this establishment stage (also called the late or hierarchical stage) has been attributed to variable reprogramming factor expression and degradation, proliferation rates, or the stochastic nature of remodeling various epigenetic marks (or barriers). Simple expression of 1-3 cell surface markers could be identified to distinguish various subpopulations within these maps, such as SSEA1 in mouse studies and TRA-1-60 in human studies. These strategies have been used to fractionate and enrich for iPSC colony identification within these heterogeneous cultures. Our regression model based on nuclear subpopulation analysis outperformed such a surface marker strategy (Figure 5), indicating multidimensional analysis can be useful for isolating desired cell types from these cultures. Given the large variety of states observed and combinations of factors or small molecules used in reprogramming, multidimensional, systems analysis is likely useful in the mapping and purification of cell states during epigenetic reprogramming.

Microenvironmental signaling from cell-cell contact, mechanotransduction, and paracrine soluble factors within the μFeatures are likely reflected in the nuclear morphology and organization. Such signaling can establish different “in vitro niches” with cell culture systems, and microenvironmental signaling is known to vary across discrete cell types described during stem cell differentiation (Liu et al., 2010) and somatic cell reprogramming (Caiazzo et al., 2016). We could utilize the nuclear morphology and organization data to distinguish cell types better than using single cell analysis (Figure 4—figure supplement 1C, **D**) which perturbs or eliminates entirely microenvironmental signaling during dissociation. Notably, clustering of cells is lost during single cell analysis, but was highly loaded in PC3 (Figure 4) and had a high coefficient in the regression model (Figure 5). Cytoskeletal staining provided little additional information for our efforts to distinguish neural progenitors, fibroblasts and iPSCs (data not shown), indicating that nuclear organization and morphology correlated with cytoskeletal changes. It is anticipated that additional stains or reporters of the cell – plasma membrane, metabolic activity, mitochondria, focal adhesion, mitotic spindle, chromatin mobility – could complement these maps and add texture to each cell state. While detailed high-content analysis of each of these aspects of cells has potential to clarify the reprogramming map, our analysis sought to identify a minimal set of subcellular characteristics to distinguish iPSCs from somatic and intermediate cell states. Because none of the measurements relied on the intensity of the nuclear stain, we anticipate a wide variety of live nuclear dyes to enable nuclear tracking to obtain these minimal measurements. These capabilities would complement other live staining strategies to mark highly dividing cells and metabolic shifts during reprogramming.

The microenvironment has recently been identified as a key regulator of reprogramming. In vivo reprogramming suggested that stress signaling within the microenvironment promotes reprogramming within a variety of tissues. Physical confinement of reprogramming cell populations to islands of a few hundreds of cells with our microcontact printed plates promoted cell fate transitions. Confinement on this length scale (100-1000 μm) also promoted reprogramming within microfluidics and microchannels (Downing et al., 2013; Luni et al., 2016). Mechanisms of extracellular accumulation of secreted factors produced by the cells already undergoing reprogramming could provide cues for other surrounding cells to progress through reprogramming. The dilution of factors can both promote or inhibit reprogramming at various stages. In mouse experiments, when dynamic culture is applied during the establishment phase of reprogramming (Sia et al., 2016), efficiency is increased due to convective mixing that prevents cell cycle arrest as cells reached confluency.

Direct changes to nuclear packing, organization and epigenetic state are also possible through mechanotransduction from the cytoskeleton to the nuclear matrix. Alignment of reprogramming cells within microchannels promoted increased global acetylation of histone 3 (Downing et al., 2013). Reprogramming on aligned nanofibers however did not result in lasting changes to nuclear shape or global acetylation state of histone 3 (Cordie et al., 2014). ‘Open’ and ‘closed’ chromatin reflect changes in global gene expression and epigenetic landscapes change during reprogramming (Hanna et al., 2010; Mattout et al., 2011). These epigenetic changes can be seen in nuclear size as stem cells have smaller nuclei than mature cells. These changes have been characterized in the reprogramming of mouse cells (Mattout et al., 2011; Smith et al., 2010), and by us in human cells (Cordie et al., 2014). Nuclear geometry is also dramatically altered during reprogramming (Cordie et al., 2014; Mattout et al., 2011). Furthermore, the packing and mobility of chromatin change during cell fate changes, notably at points during stem cell differentiation (Bickmore and van Steensel, 2013; Meshorer et al., 2006) and at the beginning and end of reprogramming (Mattout et al., 2011). Watching these changes *in situ* during stem cell differentiation permits the isolation of specific cell types of defined potency (Liu et al., 2010; Treiser et al., 2010). More recently, super-resolution imaging of nuclei has been able to forecast transitions within stem cell cultures (Kim et al., 2017). Similar studies with our microcontact printed substrates could characterize these chromatin dynamics during reprogramming, notably during the heterogeneous establishment stage of reprogramming.

Limitations of the current micropatterning approach include the spectrum of reprogramming strategies tested, 2D imaging, culture duration, and potential issues with dye compatibility for live cell imaging. First, our map and results may be restricted to the Oct4, Sox2, Klf4 and c-Myc combination in essential eight-based media, and different reprogramming cocktails (e.g., Lin28, Nanog and various small molecule cocktails) may involve different trajectories and transitions. Further, the starting cell type, fibroblasts, could be varied to watch various modes of erasure, as reprogramming trajectories for neural progenitors can skip the MET transition (Jackson et al., 2016). Second, although we did not find significantly different predictions of reprogramming when using imaging data at different z-planes (Figure 4—figure supplement 2), comprehensive 3D imaging of each μFeature could provide maps at higher resolution to further dissect the differences in morphological and clustering changes further away from the cell-substrate interface. Third, there is a limited duration of culture before cells overgrow the μFeature and potentially detach from the pattern, depending on the balance of cell-cell adhesion versus cell-substrate adhesion. For the reprogramming experiments described here, the 300 μm radius was optimized for about 20-40 day culture, but the cell seeding density or micropatterned geometry could be easily changed (Harkness et al., 2015; Sha et al., 2013) for studying other epigenetic reprogramming processes. Finally, live tracking applications may be limited for some cell lines, as imaging of live cells through uses of fluorescent dyes may cause DNA crosslinking, damage, and toxicity. Low dye concentrations and short incubation periods can potentially avoid these artifacts (Durand and Olive, 1982; Wojcik and Dobrucki, 2008; Zhao et al., 2009). To mitigate these effects further, different dyes can also be tested (Martin et al., 2005).

*In situ* analysis revealed a biphasic pattern of YAP localization within cells. The kinetics of progression through these changes is faster on micropatterns than on standard substrates. The faster kinetics are YAP-dependent, as hyperactive TAZ slowed erasure of somatic identity. YAP-independent proliferative effects could also be promoted by the micropatterns - since by day 8 of erasure, cells proliferated faster on the micropatterns despite lower YAP activity compared to unpatterned substrates. This higher proliferation rate could be due to the activity of reprogramming factors driving cell cycle progression as previously noted (Hanna et al., 2009; Kawamura et al., 2009; Marión et al., 2009). The erased, intermediate cells then become less dense, likely from apoptosis, which leads to lower cell density and recovery of YAP activity on the micropatterns during establishment. Similar decreases in cell density after erasure have been seen before (Qin et al., 2012). Higher diameter μFeatures have lower reprogramming efficiency compared to lower diameter μFeatures (Figure 4C), which likely reflects the higher diameter μFeatures inability to reduce YAP activity during erasure. For μFeature geometries such as stars and squares where local confinement may be high at corners, reprogramming efficiency was unaffected, presumably because density-dependent mechanisms to promote erasure are not acting at the single cell length scale (tens of microns), but rather long-range, across the hundreds of microns of the entire μFeature. The increases in reprogramming for μFeatures below 450 microns indicate such long-range mechanotransductive signaling mechanisms are active during micropatterning and are YAP-dependent because reprogramming efficiency increases on micropatterns were abrogated by hyperactive TAZ within the Hippo pathway.

Even with PSCs, our micropatterns had lower basal YAP activity than unpatterned substrates, presumably because confinement leads to lower cytoskeletal tension. These cells still retained all markers of pluripotent cells. This lower basal YAP activity for pluripotency on the micropatterns could also set the goal lower for the establishment phase, ultimately helping to increase the frequency of successful reprogramming. While YAP expression was seen to help in OSK reprogramming of mouse fibroblasts (Lian et al., 2010), this may be due to differences between signaling pathways used by human and mouse cells. Further, for the mouse experiments, the low proliferative drive during erasure from the absence of Myc could lead to lower cell densities, which would allow erasure to complete and allowing for YAP to assist in establishment. The levels of YAP activity during erasure in our hyperactive YAP scenario with OSKM at high cell densities were likely too high compared to these experiments and ultimately leading to slower erasure.

Fully pluripotent iPSCs were readily isolated from the micropatterns. The presence of nearly 100% pure iPSCs was surprising (Figure 6A), given the overall efficiencies of reprogramming when considering the entire well. Future work with fate mapping and clonal tracing (Lu et al., 2011; Schepers et al., 2008; Wong et al., 2005) is needed to confirm that the final cells arose from multiple cells that progressed through the establishment transition. Because all cell types firmly adhered to the surface when we performed co-culture analysis (Figure 1—figure supplement 1), we anticipate that some enhance local conversion occurred on these μFeatures. The local conversion involves differential microenvironmental signaling through the YAP/TAZ pathway and proliferation, but could also involve other mechanotransduction, cell-cell contact, and paracrine signaling mechanisms that have been linked to reprogramming. Further, the chromatin dynamics and organization could be affected from the higher cell densities and lower Hippo signaling. Utilization of the micropatterned substrates to probe these mechanisms has strong potential to provide deeper insight in the biophysical and microenvironmental signaling involved in epigenetic reprogramming. The microscale approach is complementary to other imaging modalities for dissecting complex reprogramming cultures (e.g., on cell surface markers (Paull et al., 2015; Smith et al., 2010; Tanabe et al., 2013), metabolic markers (Muthusamy et al., 2014), or dye uptake (Hirata et al., 2014; Smith et al., 2010)) and could be implemented in a variety of formats (e.g., microfluidic (Luni et al., 2016), μShear (Singh et al., 2013)). Enhanced capabilities to produce iPSCs on micropatterned substrates move both allogeneic and autologous PSC-based regenerative medicine closer to the clinic and use in precision medicine (Andrews et al., 2014; Inoue et al., 2014; Yaffe et al., 2016).

## MATERIALS AND METHODS

### Cell culture and derivation of cell lines

All reprogramming experiments were performed using the previously reported C1.2 human secondary fibroblast line (Cordie et al., 2014), which incorporates the stably integrated transgenes Oct4, Sox2, Klf4 and c-Myc on doxycycline (DXC)-inducible cassettes. All iPSCs were generated through DXC-mediated reprogramming of the C1.2 line. Transgenic iPSCs constitutively expressing H2B-mCherry and LifeAct-GFP were generated via CRISPR/Cas9 introduction of the H2B-mCherry gene (Addgene #20972) to the AAVS1 locus and lentiviral transduction of LifeAct-GFP (Addgene #22212) followed by clonal isolation of homogeneous cell lines. H2B-mCherry and LifeAct-GFP expressing fibroblasts were then differentiated from this iPSC line via embryoid body differentiation (Cordie et al., 2014; Harkness et al., 2015; Hockemeyer et al., 2008). During reprogramming studies, the TAZ S89A transgene (Addgene #52084) was introduced via lentivirus into C1.2 fibroblasts three days prior to the onset of reprogramming.

Once derived, all fibroblasts were maintained on gelatin-coated tissue culture plastic in fibroblast media containing DMEM-high glucose (Life Technologies) supplemented with 10% Fetal Bovine Serum (Life Technologies), 1 mM L-glutamine (Life Technologies), 1% non-essential amino acids (Millipore) and 1% Penicillin/Streptomycin (LifeTechnologies) and passaged with 0.05% trypsin (Life Technologies) every 3-5 days. All pluripotent cells were maintained in E8 media formulated in-house according to an established recipe (Beers et al., 2012; Chen et al., 2011) on Matrigel-coated polystyrene tissue culture plates and passaged with Versene EDTA (Life Technologies) every 3-5 days. All cells were maintained at 37°C and 5% CO_2_.

### μCP Well Plate construction

μCP Well Plates were constructed as previously described (Harkness et al., 2015). Large PDMS stamps were used with traditional microcontact printing techniques (Sha et al., 2013) to pattern a thin sheet of glass (Coresix Precision Glass, Inc.) which had been precut to the size of a standard tissue culture plate. Once patterned, the glass was then fastened to the bottom of a bottomless well plate via medical-grade double sided tape (ARcare 90106). Bottomless well plates were made in-house by removing well bottoms of standard tissue culture plates (Fisher Scientific) using a laser cutter (Universal Lasers Systems) or purchased directly (Greiner Bio-One).

### Reprogramming and isolation of iPSC lines

C1.2 secondary fibroblasts were seeded onto patterned or unpatterned surfaces in fibroblast media one day prior to reprogramming initiation. The following day, media was switched to E7 (E8 without TGF-β) supplemented with hydrocortisone (Beers et al., 2012), and DXC was added at 2 μg/mL (5 μM) to activate expression of the reprogramming factors. Media was changed every other day. To isolate pure iPSC lines, candidate colonies were picked from micropatterns using a 200 μL micropipette tip and transferred to Matrigel-coated tissue culture plastic in E8 media with DXC. If additional purification was required, one additional manual picking step with a 200 μL micropipette tip was performed. After the first passage, DXC was removed from E8 media and iPSCs were maintained in standard pluripotent culture as described above. During picking and subsequent passaging, the culture media was often supplemented with the Rho kinase inhibitor Y-27632 (Sigma) to encourage cell survival and establish clonal lines.

### Antibodies and Staining

All cells were fixed for 15 minutes with 4% paraformaldehyde in PBS (Sigma) and permeabilized with 0.5% Triton-X (Sigma) for >4 hours before staining. Hoechst (Life Technologies H1399) was used at 5 μg/mL to stain nuclei. Primary antibodies were applied overnight in a blocking buffer of 5% donkey serum (Sigma) at the following concentrations: CD44-PE (BD Biosciences 555479), 1:200; TRA-1-60 (Millipore MAB4360) 1:100; Nanog (R&D Systems AF1997) 1:200; Oct4 (BD Biosciences 560186) 1:10; Pax6 (DSHB AB_528427) 1:200; YAP (Santa Cruz sc-101199) 1:300. Secondary antibodies were obtained from Life Technologies and applied in a blocking buffer of 5% donkey serum for one hour at concentrations of 1:400 – 1:800. Phalloidin-TRITC was used at 10 nM to visualize F-actin. For live-cell stains, antibodies were added to the appropriate cell culture media at equivalent dilutions to those used for fixed cells. Two-hour incubations were used for primary antibodies, followed by 30-minute incubations for secondary antibodies.

### High-content analysis

High-content image analysis was performed similarly to previously published methods (Harkness et al., 2015). A Nikon Eclipse-Ti epifluorescence microscope was used to acquire single 10x images of each micropattern, and a Nikon AR1 confocal microscope was used to acquire 60x stitched images of each micropattern using the z-plane closest to the glass substrate for reprogramming studies. Images were fed directly into the image analysis software CellProfiler (Carpenter et al., 2006), which analyzed images as described in Figure 3—figure supplement 1. Objects <300 pixels in area were filtered out of the data set to exclude apoptotic or other debris, and neighbors were identified by expanding objects until all pixels on the object boundaries were touching one another. Two objects are neighbors if any of their boundary pixels are adjacent after expansion. Measurement of YAP staining intensity across a micropattern was obtained from the *MeanFrac* Radial Distribution function, which measures the fraction of total intensity normalized by the fraction of pixels at a given radius.

### PCA and PLSR

CellProfiler measurements were averaged across each μFeature and fed as mean values into PCA and PLSR analysis via MATLAB software. PLSR was performed using the NIPALS method and predictions were tested and validated using K-Fold validation with 7 folds. Actual Nanog expression levels were obtained by manually tracing the areas of each micropattern expressing Nanog and dividing by the area stained by Hoechst.

### Statistics

All error bars and box plot notches are presented as 95% confidence intervals. *p*-values were calculated using unpaired two-tailed Student’s *t*-tests with unequal variance or one-way ANOVA using MATLAB. Statistical tests were deemed significant at α≤0.05. Unless directly stated, all exact *p*-values can be found in supplementary tables. Technical replicates are defined as distinct μFeatures within an experiment. Biological replicates are experiments performed at distinct time points during reprogramming. Outliers were identified as 1.5*IQR and excluded from statistical test. Outlier data points are still shown for illustration. No *a priori* power calculations were performed.

## Acknowledgments

We thank members of the Saha lab for helpful discussion and comments on the manuscript. We also thank plasmid depositors to Addgene.

## Author contributions

T.H, J.C-S., S.S. planned research and analyzed data. T.H, K.S. designed experiments. T.H., R.P. performed experiments and J.C-S, T.H wrote the manuscript with input from all authors. K.S. supervised research.

## SUPPLEMENTARY FIGURES

**Figure 1—figure supplement 1.**
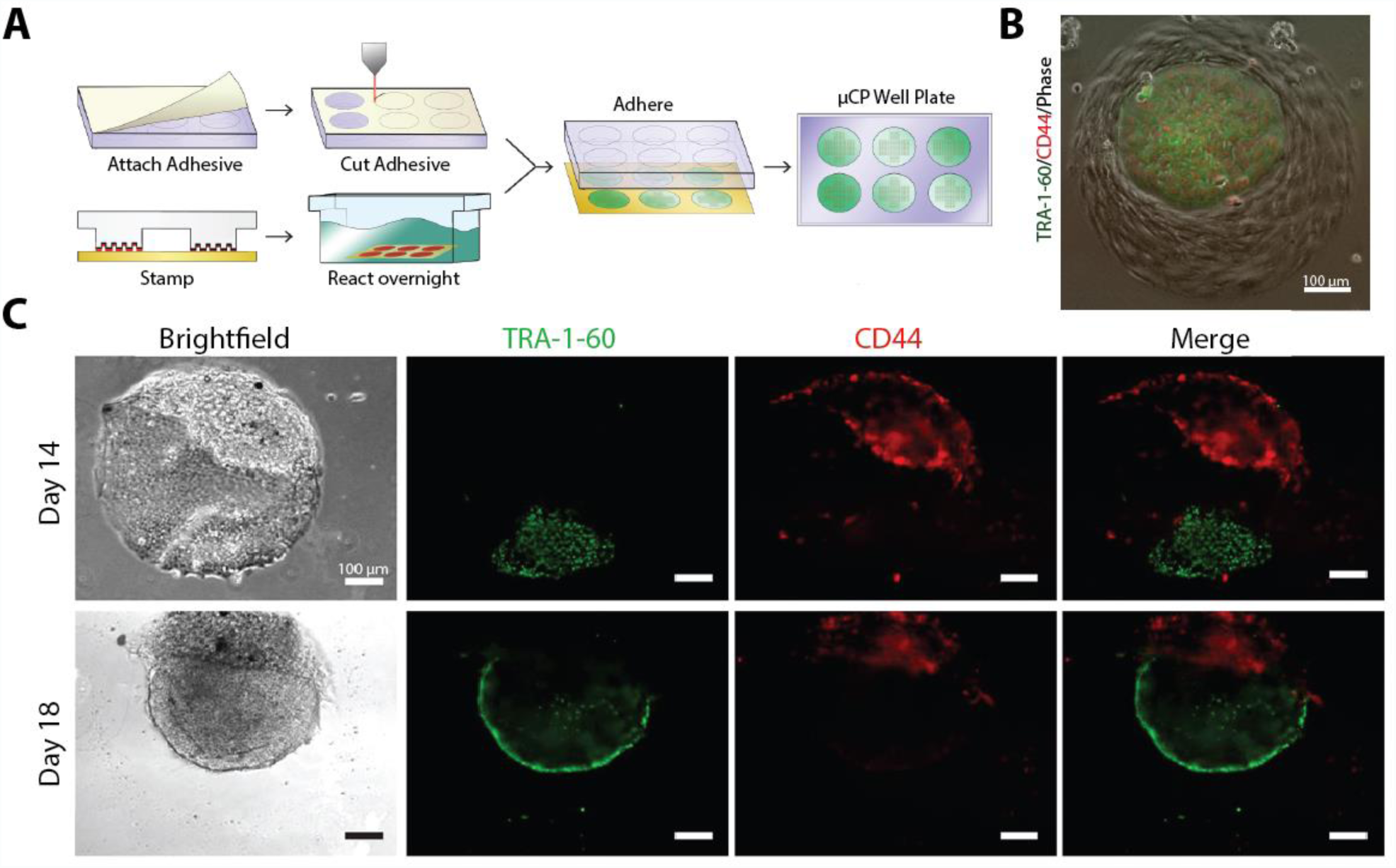
Micropatterned plates enable live tracking of reprogramming. **A)** Schematic showing the creation of μCP Well Plates. Gold-coated glass is stamped with initiator solution using PDMS mold and then reacted in PEG solution. Glass is then combined with a standard well plate that has had the bottom cut out and a double side adhesive attached. **B)** Coculture of two different cell types on a single μFeature. Brightfield only cells are fibroblasts. H2B-mCherry, Actin-GFP cells are PSCs. **C)** Representative images of the progression of a cell aggregate on a single μFeature through establishment. Cells were live stained using antibodies against TRA-1-60 (green) and CD44 (red).

**Figure 2—figure supplement 1.**
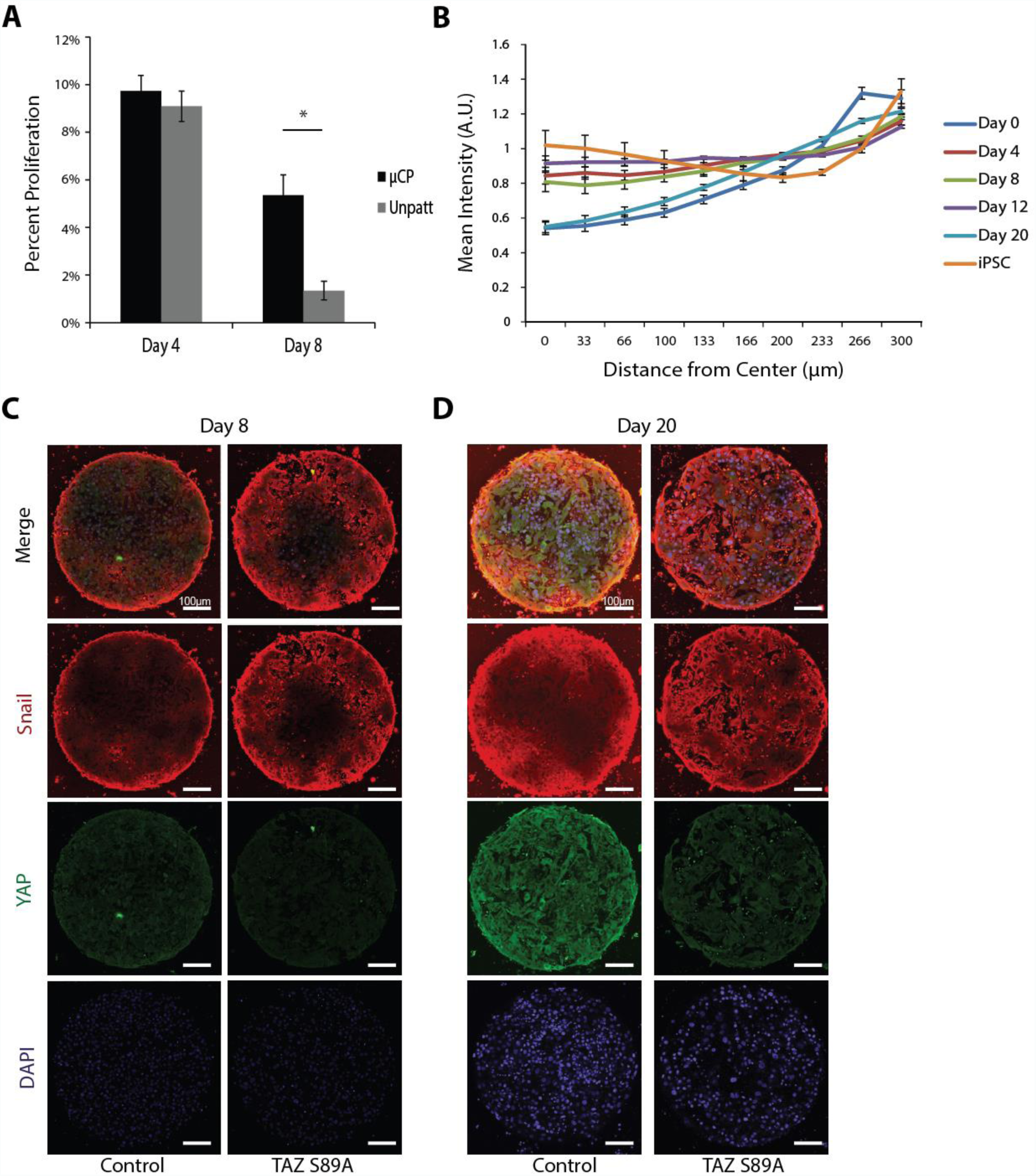
Proliferation, YAP and Snail levels within cells undergoing reprogramming. **A)** Proliferation rates during early reprogramming on μFeatures compared to unpatterned substrates. Over 3-fold as many cells are actively proliferating, on day 8 as compared to unpatterned. Data quantified using Click-iT EdU (n=20 technical replicates Student’s two-tailed t-test *p*=0.18, 8.4*10^−9^). **B)** YAP intensity as function of distance from center of μFeature. Intensity increased on the edge of the μFeature due to mechanical strain. **C, D)** Representative images of YAP and Snail dynamics during erasure using hyperactive TAZ on Day 8 (C) and Day 20 (D)

**Figure 3—figure supplement 1.**
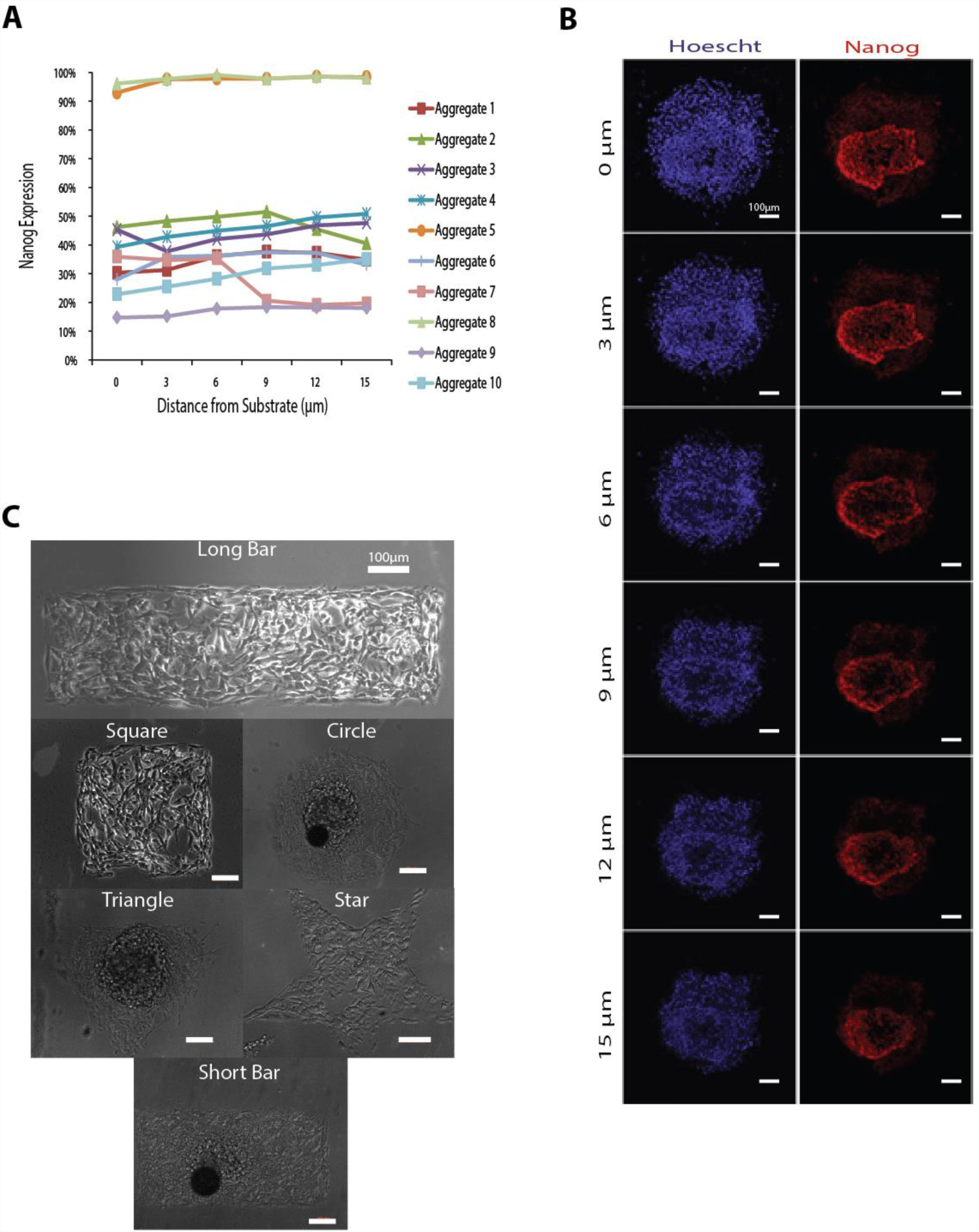
Z-stack analysis and representative micropattern geometries. **A)** Nanog expression as a function of z-distance from the substrate. Expression levels did not vary with height. **B)** Representative images of Nanog expression with increasing z-distance. **C)** Representative images of reprogramming cells on five different μFeature shapes.

**Figure 4—figure supplement 1.**
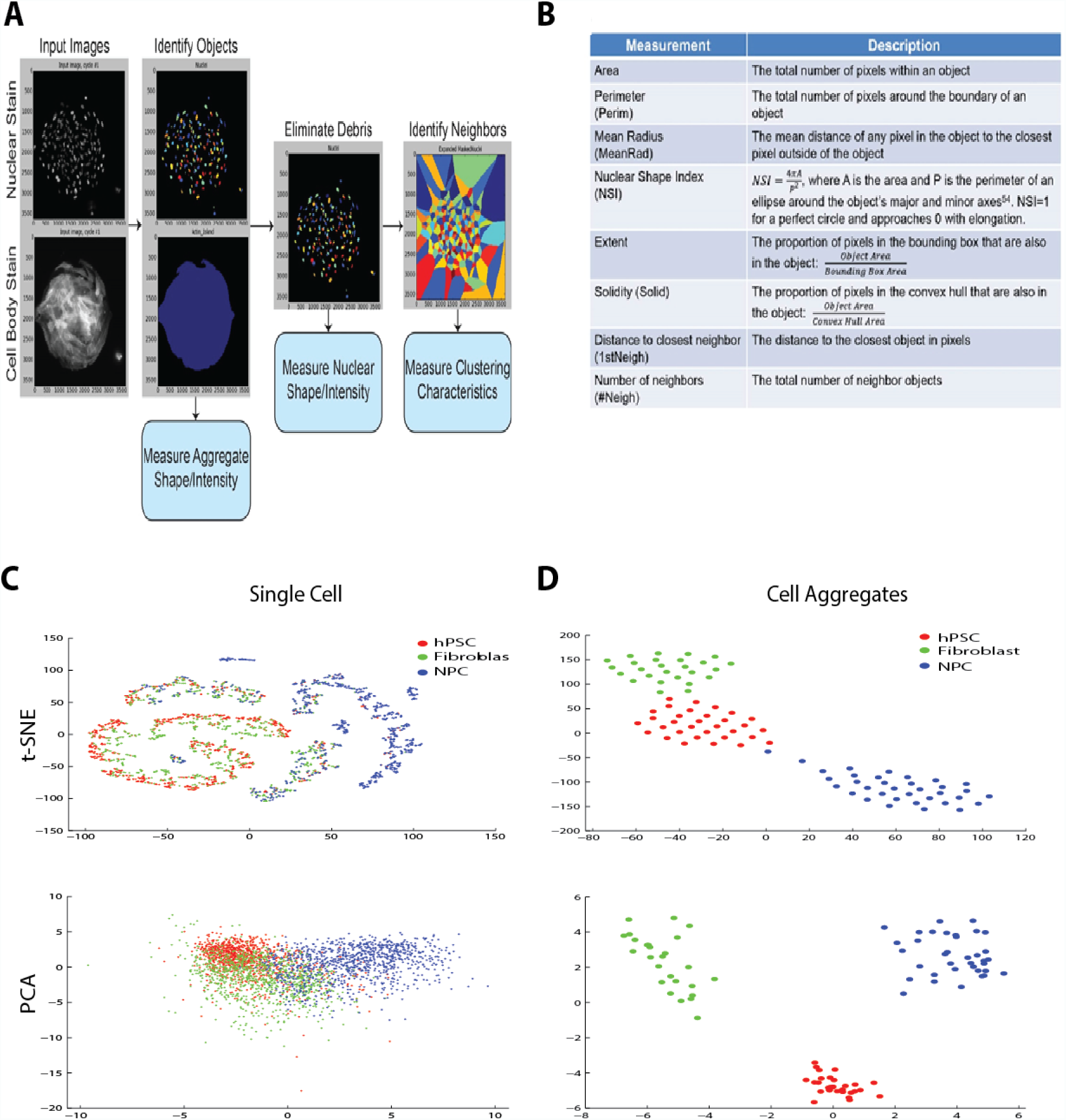
Nuclear characteristics within μFeatures can separate cell types better than single cell nuclear characteristics. **A)** CellProfiler pipeline to identify nuclear characteristics. **B)** Eight core nuclear characteristics used to create reprogramming models. **C)** Separation of cell types using single cell characteristics with t-SNE (top) and PCA (bottom) algorithms. No clusters were identified. **D)** Separation of cell types using nuclear characteristics classified averaged over μFeatures. Distinct cell populations were visible by eye using t-SNE and PCA.

**Figure 4—figure supplement 2.**
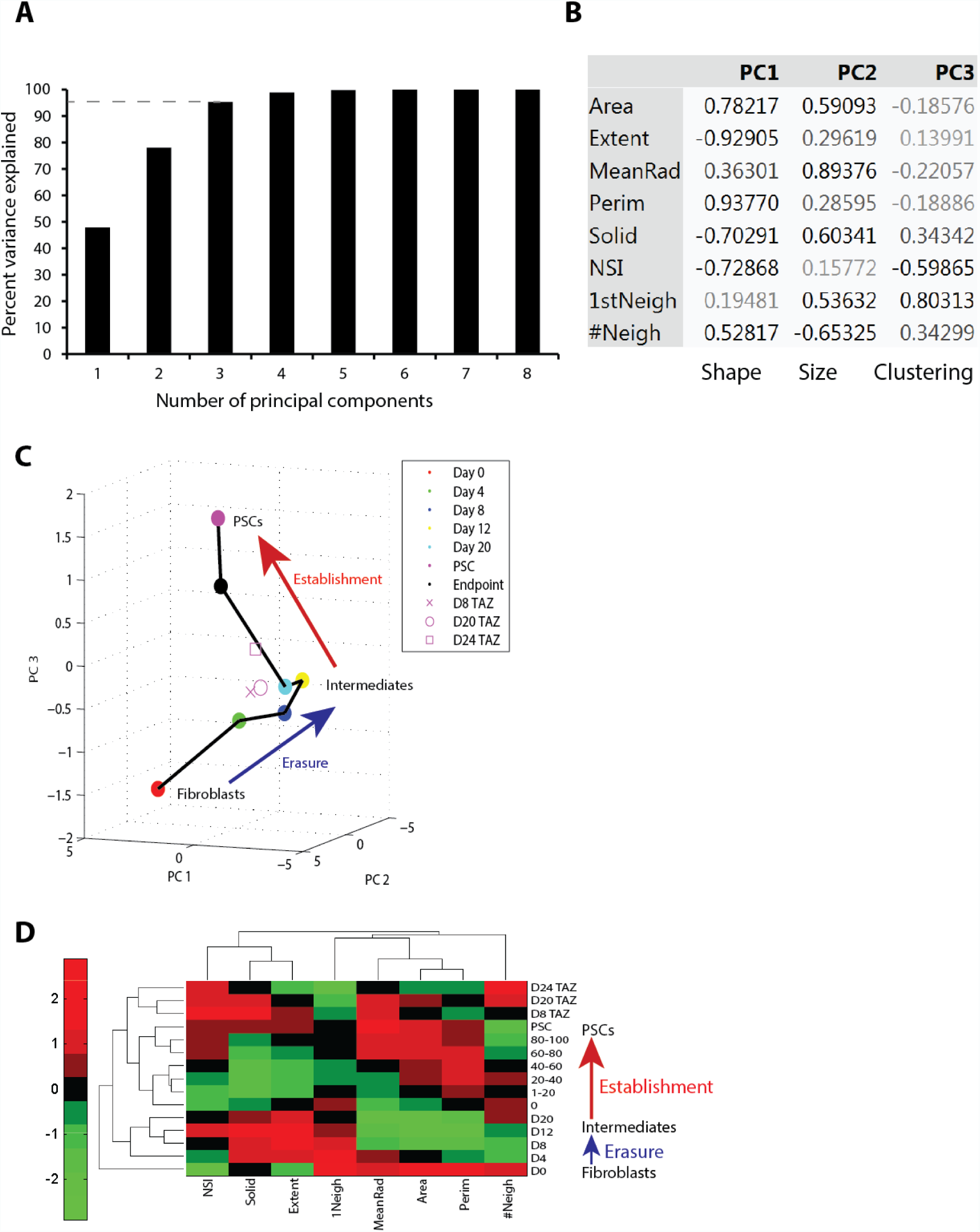
Principal components analysis and clustering analysis of nuclear characteristics. **A)** Three PCs explained 95% of the variance in the PCA model while additional components had diminishing returns. **B)** Loadings of PCA model for each of the first three PCs. PCs can be broken down to correspond to shape, size, and clustering of the nuclei. **C)** Reprogramming spiral including μFeatures transduced with hyperactive TAZ. D8 and D20 μFeatures clustered closely to D4-8 of standard reprogramming. **D)** Hierarchical clustering of reprogramming including hyperactive TAZ μFeatures. These μFeatures clustered close to each other.

**Figure 6—figure supplement 1.**
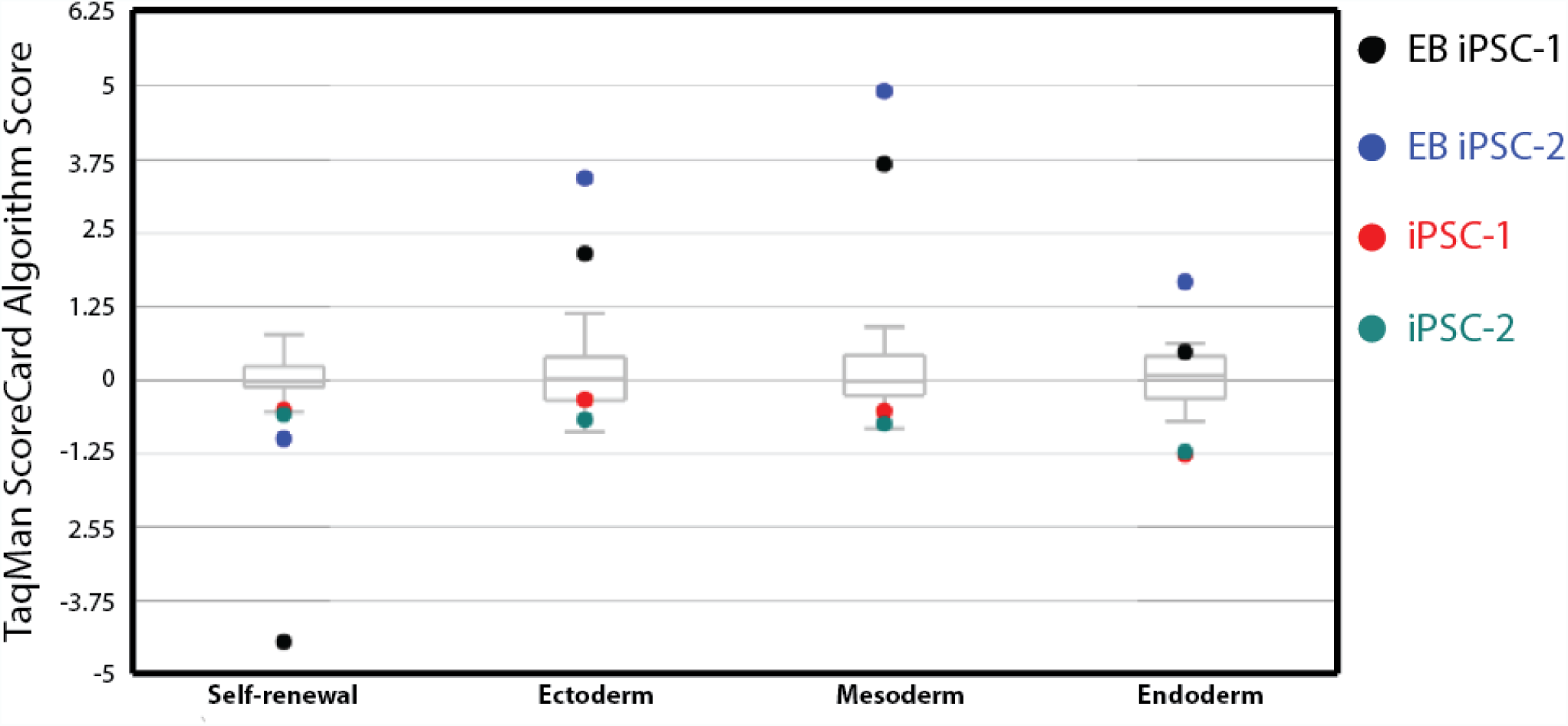
Standardized gene expression from ScoreCard™ assay. Both iPSC clones upregulated genes associated with self-renewal. Embryoid bodies formed from both clones expressed genes associated with all three germ layers. Data is compared to 9 standard hPSC lines (grey box-and-whisker plots).

**Figure 1—Source data 1.** Cell density on μFeatures and unpatterened substrates during reprogramming.

**Figure 1—Source data 2.** YAP localization on μFeatures and unpatterened substrates during reprogramming.

**Figure 2—Source data 1.** Cell proliferation during reprogramming on day 4 and day 8.

**Figure 2—Source data 2.** Cell density, YAP localization and intensity, and Snail intensity during reprogramming in wild-type and hTAZ infected cells.

**Figure 4—Source data 1.** Nuclear characteristics of reprogramming cells at 6 time points used to form PCA model.

**Supplementary File:** Statistical breakdown of larger data sets.

## REFERENCES

Andrews, P.W., Cavagnaro, J., Cavanagro, J., Deans, R., Feigal, E., Feigel, E., Horowitz, E., Keating, A., Rao, M., Turner, M., et al. (2014). Harmonizing standards for producing clinical-grade therapies from pluripotent stem cells. Nat. Biotechnol. 32, 724–726.

Aragona, M.A.-P., Tito AU-Manfrin, Andrea AU-Giulitti, Stefano AU-Michielin, Federica AU- Elvassore, Nicola AU-Dupont, Sirio AU-Piccolo, Stefano (2013). A mechanical checkpoint controls multicellular growth through YAP/TAZ regulation by actin-processing factors. Cell 154, 1047–1059.

Bauwens, C.L., Peerani, R., Niebruegge, S., Woodhouse, K.A., Kumacheva, E., Husain, M., and Zandstra, P.W. (2008). Control of Human Embryonic Stem Cell Colony and Aggregate Size Heterogeneity Influences Differentiation Trajectories. STEM CELLS 26, 2300–2310.

Beers, J., Gulbranson, D.R., George, N., Siniscalchi, L.I., Jones, J., Thomson, J.A., and Chen, G. (2012). Passaging and colony expansion of human pluripotent stem cells by enzyme-free dissociation in chemically defined culture conditions. Nat. Protoc. 7, 2029–2040.

Bickmore, W.A., and van Steensel, B. (2013). Genome architecture: domain organization of interphase chromosomes. Cell 152, 1270–1284.

Bratt-Leal, A.M.A.-C., Richard L. AU-McDevitt, Todd C. (2009). Engineering the embryoid body microenvironment to direct embryonic stem cell differentiation. Biotechnol. Prog. 25, 43–51.

Bronshtein, I., Kepten, E., Kanter, I., Berezin, S., Lindner, M., Redwood, A.B., Mai, S., Gonzalo, S., Foisner, R., and Shav-Tal, Y. (2015). Loss of lamin A function increases chromatin dynamics in the nuclear interior. Nat. Commun. 6.

Buganim, Y., Faddah, D.A., Cheng, A.W., Itskovich, E., Markoulaki, S., Ganz, K., Klemm, S.L., van Oudenaarden, A., and Jaenisch, R. (2012). Single-cell expression analyses during cellular reprogramming reveal an early stochastic and a late hierarchic phase. Cell 150, 1209–1222.

Cahan, P., and Daley, G.Q. (2013). Origins and implications of pluripotent stem cell variability and heterogeneity. Nat. Rev. Mol. Cell Biol. 14, 357–368.

Caiazzo, M., Okawa, Y., Ranga, A., Piersigilli, A., Tabata, Y., and Lutolf, M.P. (2016). Defined three-dimensional microenvironments boost induction of pluripotency. Nat. Mater. 15, 344–352.

Carlson-Stevermer, J., Goedland, M., Steyer, B., Movaghar, A., Lou, M., Kohlenberg, L., Prestil, R., and Saha, K. (2016). High-Content Analysis of CRISPR-Cas9 Gene-Edited Human Embryonic Stem Cells. STEM Cell Rep. 6, 109–120.

Carpenter, A.E., Jones, T.R., Lamprecht, M.R., Clarke, C., Kang, I.H., Friman, O., Guertin, D.A., Chang, J.H., Lindquist, R.A., Moffat, J., et al. (2006). CellProfiler: image analysis software for identifying and quantifying cell phenotypes. Genome Biol. 7, R100.

Chan, E.M.A.-R., Sutheera AU-Park, In-Hyun AU-Manos, Philip D. AU-Loh, Yuin-Han AU-Huo, Hongguang AU-Miller, Justine D. AU-Hartung, Odelya AU-Rho, Junsung AU-Ince, Tan A. (2009). Live cell imaging distinguishes bona fide human iPS cells from partially reprogrammed cells. Nat. Biotechnol. 27, 1033–1037.

Chen, G., Gulbranson, D.R., Hou, Z., Bolin, J.M., Ruotti, V., Probasco, M.D., Smuga-Otto, K., Howden, S.E., Diol, N.R., Propson, N.E., et al. (2011). Chemically defined conditions for human iPSC derivation and culture. Nat. Methods 8, 424–429.

Choi, B., Park, K.-S., Kim, J.-H., Ko, K.-W., Kim, J.-S., Han, D.K., and Lee, S.-H. (2016). Stiffness of Hydrogels Regulates Cellular Reprogramming Efficiency Through Mesenchymal-to-Epithelial Transition and Stemness Markers. Macromol. Biosci. 16, 199–206.

Cordie, T., Harkness, T., Jing, X., Carlson-Stevermer, J., Mi, H.-Y., Turng, L.-S., and Saha, K. (2014). Nanofibrous Electrospun Polymers for Reprogramming Human Cells. Cell. Mol. Bioeng. 7, 379–393.

Dingal, P.D.P., and Discher, D.E. (2014). Material control of stem cell differentiation: challenges in nano-characterization. Curr. Opin. Biotechnol. 28, 46–50.

Downing, T.L., Soto, J., Morez, C., Houssin, T., Fritz, A., Yuan, F., Chu, J., Patel, S., Schaffer, D.V., and Li, S. (2013). Biophysical regulation of epigenetic state and cell reprogramming. Nat. Mater. 12, 1154–1162.

Dupont, S., Morsut, L., Aragona, M., Enzo, E., Giulitti, S., Cordenonsi, M., Zanconato, F., Le Digabel, J., Forcato, M., Bicciato, S., et al. (2011). Role of YAP/TAZ in mechanotransduction. Nature 474, 179–183.

Durand, R.E., and Olive, P.L. (1982). Cytotoxicity, Mutagenicity and DNA damage by Hoechst 33342. J. Histochem. Cytochem. Off. J. Histochem. Soc. 30, 111–116.

Gingold, J.A., Fidalgo, M., Guallar, D., Lau, Z., Sun, Z., Zhou, H., Faiola, F., Huang, X., Lee, D.- F., Waghray, A., et al. (2014). A genome-wide RNAi screen identifies opposing functions of Snai1 and Snai2 on the Nanog dependency in reprogramming. Mol. Cell 56, 140–152.

Guelen, L., Pagie, L., Brasset, E., Meuleman, W., Faza, M.B., Talhout, W., Eussen, B.H., de Klein, A., Wessels, L., and de Laat, W. (2008). Domain organization of human chromosomes revealed by mapping of nuclear lamina interactions. Nature 453, 948–951.

Hanna, J., Saha, K., Pando, B., van Zon, J., Lengner, C.J., Creyghton, M.P., van Oudenaarden, A., and Jaenisch, R. (2009). Direct cell reprogramming is a stochastic process amenable to acceleration. Nature 462, 595–601.

Hanna, J.H., Saha, K., and Jaenisch, R. (2010). Pluripotency and cellular reprogramming: facts, hypotheses, unresolved issues. Cell 143, 508–525.

Hansson, J., Rafiee, M.R., Reiland, S., Polo, J.M., Gehring, J., Okawa, S., Huber, W., Hochedlinger, K., and Krijgsveld, J. (2012). Highly Coordinated Proteome Dynamics during Reprogramming of Somatic Cells to Pluripotency. Cell Rep. 2, 1579–1592.

Harkness, T., McNulty, J.D., Prestil, R., Seymour, S.K., Klann, T., Murrell, M., Ashton, R.S., and Saha, K. (2015). High-content imaging with micropatterned multiwell plates reveals influence of cell geometry and cytoskeleton on chromatin dynamics. Biotechnol. J. 10, 1555–1567.

Hendrix, M.J.C., Seftor, E.A., Seftor, R.E.B., Kasemeier-Kulesa, J., Kulesa, P.M., and Postovit, L.-M. (2007). Reprogramming metastatic tumour cells with embryonic microenvironments. Nat. Rev. Cancer 7, 246–255.

Hirata, N., Nakagawa, M., Fujibayashi, Y., Yamauchi, K., Murata, A., Minami, I., Tomioka, M., Kondo, T., Kuo, T.-F., Endo, H., et al. (2014). A Chemical Probe that Labels Human Pluripotent Stem Cells. Cell Rep. 6, 1165–1174.

Hockemeyer, D., Soldner, F., Cook, E.G., Gao, Q., Mitalipova, M., and Jaenisch, R. (2008). A Drug-Inducible System for Direct Reprogramming of Human Somatic Cells to Pluripotency. Cell Stem Cell 3, 346–353.

Hsiao, C., Lampe, M., Nillasithanukroh, S., Han, W., Lian, X., and Palecek, S.P. (2016). Human pluripotent stem cell culture density modulates YAP signaling. Biotechnol. J. n/a – n/a.

Inoue, H., Nagata, N., Kurokawa, H., and Yamanaka, S. (2014). iPS cells: a game changer for future medicine. EMBO J. 33, 409–417.

Iskratsch, T., Wolfenson, H., and Sheetz, M.P. (2014). Appreciating force and shape—the rise of mechanotransduction in cell biology. Nat. Rev. Mol. Cell Biol. 15, 825–833.

Jackson, S.A., Olufs, Z.P.G., Tran, K.A., Zaidan, N.Z., and Sridharan, R. (2016). Alternative Routes to Induced Pluripotent Stem Cells Revealed by Reprogramming of the Neural Lineage. STEM Cell Rep. 6, 302–311.

Jain, N., Iyer, K.V., Kumar, A., and Shivashankar, G.V. (2013). Cell geometric constraints induce modular gene-expression patterns via redistribution of HDAC3 regulated by actomyosin contractility. Proc. Natl. Acad. Sci. 110, 11349–11354.

Kawamura, T., Suzuki, J., Wang, Y.V., Menendez, S., Morera, L.B., Raya, A., Wahl, G.M., and Belmonte, J.C.I. (2009). Linking the p53 tumour suppressor pathway to somatic cell reprogramming. Nature 460, 1140–1144.

Kim, J.J., Bennett, N.K., Devita, M.S., Chahar, S., Viswanath, S., Lee, E.A., Jung, G., Shao, P.P., Childers, E.P., Liu, S., et al. (2017). Optical High Content Nanoscopy of Epigenetic Marks Decodes Phenotypic Divergence in Stem Cells. Sci. Rep. 7, 39406.

Kind, J., Pagie, L., de Vries, S.S., Nahidiazar, L., Dey, S.S., Bienko, M., Zhan, Y., Lajoie, B., de Graaf, C.A., and Amendola, M. (2015). Genome-wide maps of nuclear lamina interactions in single human cells. Cell 163, 134–147.

Koche, R.P., Smith, Z.D., Adli, M., Gu, H., Ku, M., Gnirke, A., Bernstein, B.E., and Meissner, A. (2011). Reprogramming factor expression initiates widespread targeted chromatin remodeling. Cell Stem Cell 8, 96–105.

Kreiner, T., and Irion, S. (2013). Whole-Genome Analysis, Stem Cell Research, and the Future of Biobanks. Cell Stem Cell 12, 513–516.

Lee, S.E., Kamm, R.D., and Mofrad, M.R. (2007). Force-induced activation of talin and its possible role in focal adhesion mechanotransduction. J. Biomech. 40, 2096–2106.

Li, R., Liang, J., Ni, S., Zhou, T., Qing, X., Li, H., He, W., Chen, J., Li, F., Zhuang, Q., et al. (2010a). A mesenchymal-to-epithelial transition initiates and is required for the nuclear reprogramming of mouse fibroblasts. Cell Stem Cell 7, 51–63.

Li, Z., Zhao, B., Wang, P., Chen, F., Dong, Z., Yang, H., Guan, K.-L., and Xu, Y. (2010b). Structural insights into the YAP and TEAD complex. Genes Dev. 24, 235–240.

Lian, I., Kim, J., Okazawa, H., Zhao, J., Zhao, B., Yu, J., Chinnaiyan, A., Israel, M.A., Goldstein, L.S.B., Abujarour, R., et al. (2010). The role of YAP transcription coactivator in regulating stem cell self-renewal and differentiation. Genes Dev. 24, 1106–1118.

Liao, B., Bao, X., Liu, L., Feng, S., Zovoilis, A., Liu, W., Xue, Y., Cai, J., Guo, X., Qin, B., et al. (2011). MicroRNA cluster 302-367 enhances somatic cell reprogramming by accelerating a mesenchymal-to-epithelial transition. J. Biol. Chem. 286, 17359–17364.

Liu, E., Gordonov, S., Treiser, M.D., and Moghe, P.V. (2010). Parsing the early cytoskeletal and nuclear organizational cues that demarcate stem cell lineages. Cell Cycle 9, 2108–2117.

Liu-Chittenden, Y., Huang, B., Shim, J.S., Chen, Q., Lee, S.-J., Anders, R.A., Liu, J.O., and Pan, D. (2012). Genetic and pharmacological disruption of the TEAD–YAP complex suppresses the oncogenic activity of YAP. Genes Dev. 26, 1300–1305.

Low, B.C., Pan, C.Q., Shivashankar, G., Bershadsky, A., Sudol, M., and Sheetz, M. (2014). YAP/TAZ as mechanosensors and mechanotransducers in regulating organ size and tumor growth. FEBS Lett. 588, 2663–2670.

Lu, R., Neff, N.F., Quake, S.R., and Weissman, I.L. (2011). Tracking single hematopoietic stem cells in vivo using high-throughput sequencing in conjunction with viral genetic barcoding. Nat. Biotechnol. 29, 928–933.

Luni, C., Giulitti, S., Serena, E., Ferrari, L., Zambon, A., Gagliano, O., Giobbe, G.G., Michielin, F., Knöbel, S., Bosio, A., et al. (2016). High-efficiency cellular reprogramming with microfluidics. Nat. Methods 13, 446–452.

Maaten, L. van der, and Hinton, G. (2008). Visualizing Data using t-SNE. J. Mach. Learn. Res. 9, 2579–2605.

Marión, R.M., Strati, K., Li, H., Murga, M., Blanco, R., Ortega, S., Fernandez-Capetillo, O., Serrano, M., and Blasco, M.A. (2009). A p53-mediated DNA damage response limits reprogramming to ensure iPS cell genomic integrity. Nature 460, 1149–1153.

Martin, R.M., Leonhardt, H., and Cardoso, M.C. (2005). DNA labeling in living cells. Cytom. Part J. Int. Soc. Anal. Cytol. 67, 45–52.

Mattout, A., Biran, A., and Meshorer, E. (2011). Global epigenetic changes during somatic cell reprogramming to iPS cells. J. Mol. Cell Biol. 3, 341–350.

Meshorer, E., Yellajoshula, D., George, E., Scambler, P.J., Brown, D.T., and Misteli, T. (2006). Hyperdynamic Plasticity of Chromatin Proteins in Pluripotent Embryonic Stem Cells. Dev. Cell 10, 105–116.

Mikkelsen, T.S., Hanna, J., Zhang, X., Ku, M., Wernig, M., Schorderet, P., Bernstein, B.E., Jaenisch, R., Lander, E.S., and Meissner, A. (2008). Dissecting direct reprogramming through integrative genomic analysis. Nature 454, 49–55.

Mitra, S.K., Hanson, D.A., and Schlaepfer, D.D. (2005). Focal adhesion kinase: in command and control of cell motility. Nat. Rev. Mol. Cell Biol. 6, 56–68.

Musah, S., Wrighton, P.J., Zaltsman, Y., Zhong, X., Zorn, S., Parlato, M.B., Hsiao, C., Palecek, S.P., Chang, Q., and Murphy, W.L. (2014). Substratum-induced differentiation of human pluripotent stem cells reveals the coactivator YAP is a potent regulator of neuronal specification. Proc. Natl. Acad. Sci. 111, 13805–13810.

Muthusamy, T., Mukherjee, O., Menon, R., Megha, P.B., and Panicker, M.M. (2014). A method to identify and isolate pluripotent human stem cells and mouse epiblast stem cells using lipid body-associated retinyl ester fluorescence. STEM Cell Rep. 3, 169–184.

Paull, D., Sevilla, A., Zhou, H., Hahn, A.K., Kim, H., Napolitano, C., Tsankov, A., Shang, L., Krumholz, K., Jagadeesan, P., et al. (2015). Automated, high-throughput derivation, characterization and differentiation of induced pluripotent stem cells. Nat. Methods 12, 885–892.

Peric-Hupkes, D., Meuleman, W., Pagie, L., Bruggeman, S.W.M., Solovei, I., Brugman, W., Gräf, S., Flicek, P., Kerkhoven, R.M., van Lohuizen, M., et al. (2010). Molecular Maps of the Reorganization of Genome-Nuclear Lamina Interactions during Differentiation. Mol. Cell 38, 603–613.

Polo, J.M., Anderssen, E., Walsh, R.M., Schwarz, B.A., Nefzger, C.M., Lim, S.M., Borkent, M., Apostolou, E., Alaei, S., Cloutier, J., et al. (2012). A Molecular Roadmap of Reprogramming Somatic Cells into iPS Cells. Cell 151, 1617–1632.

Qin, H., Blaschke, K., Wei, G., Ohi, Y., Blouin, L., Qi, Z., Yu, J., Yeh, R.-F., Hebrok, M., and Ramalho-Santos, M. (2012). Transcriptional analysis of pluripotency reveals the Hippo pathway as a barrier to reprogramming. Hum. Mol. Genet. 21, 2054–2067.

Quintanilla, R.H., Jr., Asprer, J.S.T., Vaz, C., Tanavde, V., and Lakshmipathy, U. (2014). CD44 Is a Negative Cell Surface Marker for Pluripotent Stem Cell Identification during Human Fibroblast Reprogramming. PLoS ONE 9.

Sakurai, K., Talukdar, I., Patil, V.S., Dang, J., Li, Z., Chang, K.-Y., Lu, C.-C., Delorme-Walker, V., DerMardirossian, C., and Anderson, K. (2014). Kinome-wide functional analysis highlights the role of cytoskeletal remodeling in somatic cell reprogramming. Cell Stem Cell 14, 523–534.

Samavarchi-Tehrani, P., Golipour, A., David, L., Sung, H.-K., Beyer, T.A., Datti, A., Woltjen, K., Nagy, A., and Wrana, J.L. (2010). Functional genomics reveals a BMP-driven mesenchymal-to-epithelial transition in the initiation of somatic cell reprogramming. Cell Stem Cell 7, 64–77.

Schepers, K., Swart, E., Heijst, J.W.J. van, Gerlach, C., Castrucci, M., Sie, D., Heimerikx, M., Velds, A., Kerkhoven, R.M., Arens, R., et al. (2008). Dissecting T cell lineage relationships by cellular barcoding. J. Exp. Med. 205, 2309–2318.

Sha, J., Lippmann, E.S., McNulty, J., Ma, Y., and Ashton, R.S. (2013). Sequential Nucleophilic Substitutions Permit Orthogonal Click Functionalization of Multicomponent PEG Brushes. Biomacromolecules 14, 3294–3303.

Sia, J., Sun, R., Chu, J., and Li, S. (2016). Dynamic culture improves cell reprogramming efficiency. Biomaterials 92, 36–45.

Singh, A., Suri, S., Lee, T., Chilton, J.M., Cooke, M.T., Chen, W., Fu, J., Stice, S.L., Lu, H., McDevitt, T.C., et al. (2013). Adhesion strength-based, label-free isolation of human pluripotent stem cells. Nat. Methods 10, 438–444.

Smith, Z.D., Nachman, I., Regev, A., and Meissner, A. (2010). Dynamic single-cell imaging of direct reprogramming reveals an early specifying event. Nat. Biotechnol. 28, 521–526.

Sorrentino, G., Ruggeri, N., Specchia, V., Cordenonsi, M., Mano, M., Dupont, S., Manfrin, A., Ingallina, E., Sommaggio, R., Piazza, S., et al. (2014). Metabolic control of YAP and TAZ by the mevalonate pathway. Nat. Cell Biol. 16, 357–366.

Sun, Y., Yong, K.M.A., Villa-Diaz, L.G., Zhang, X., Chen, W., Philson, R., Weng, S., Xu, H., Krebsbach, P.H., and Fu, J. (2014). Hippo/YAP-mediated rigidity-dependent motor neuron differentiation of human pluripotent stem cells. Nat. Mater. 13, 599–604.

Tanabe, K., Nakamura, M., Narita, M., Takahashi, K., and Yamanaka, S. (2013). Maturation, not initiation, is the major roadblock during reprogramming toward pluripotency from human fibroblasts. Proc. Natl. Acad. Sci. U. S. A. 110, 12172–12179.

Toh, K.C., Ramdas, N.M., and Shivashankar, G. (2015). Actin cytoskeleton differentially alters the dynamics of lamin A, HP1α and H2B core histone proteins to remodel chromatin condensation state in living cells. Integr. Biol. 7, 1309–1317.

Treiser, M.D., Yang, E.H., Gordonov, S., Cohen, D.M., Androulakis, I.P., Kohn, J., Chen, C.S., and Moghe, P.V. (2010). Cytoskeleton-based forecasting of stem cell lineage fates. Proc. Natl. Acad. Sci. 107, 610–615.

Tsankov, A.M., Akopian, V., Pop, R., Chetty, S., Gifford, C.A., Daheron, L., Tsankova, N.M., and Meissner, A. (2015). A qPCR ScoreCard quantifies the differentiation potential of human pluripotent stem cells. Nat. Biotechnol. 33, 1182–1192.

Varelas, X., Sakuma, R., Samavarchi-Tehrani, P., Peerani, R., Rao, B.M., Dembowy, J., Yaffe, M.B., Zandstra, P.W., and Wrana, J.L. (2008). TAZ controls Smad nucleocytoplasmic shuttling and regulates human embryonic stem-cell self-renewal. Nat. Cell Biol. 10, 837–848.

Vogel, V., and Sheetz, M. (2006). Local force and geometry sensing regulate cell functions. Nat. Rev. Mol. Cell Biol. 7, 265–275.

Wheeler, B.C., Corey, J.M., Brewer, G.J., and Branch, D.W. (1999). Microcontact Printing for Precise Control of Nerve Cell Growth in Culture. J. Biomech. Eng. 121, 73–78.

Wojcik, K., and Dobrucki, J.W. (2008). Interaction of a DNA intercalator DRAQ5, and a minor groove binder SYTO17, with chromatin in live cells-influence on chromatin organization and histone-DNA interactions. Cytom. Part J. Int. Soc. Anal. Cytol. 73, 555–562.

Woltjen, K., and Stanford, W.L. (2009). Inhibition of Tgf-β Signaling Improves Mouse Fibroblast Reprogramming. Cell Stem Cell 5, 457–458.

Wong, E.T., Kolman, J.L., Li, Y.-C., Larry D. Mesner, Hillen, W., Berens, C., and Wahl, G.M. (2005). Reproducible doxycycline-inducible transgene expression at specific loci generated by Cre-recombinase mediated cassette exchange. Nucleic Acids Res. 33, e147–e147.

Xie, Q., Chen, J., Feng, H., Peng, S., Adams, U., Bai, Y., Huang, L., Li, J., Huang, J., and Meng, S. (2013). YAP/TEAD–Mediated Transcription Controls Cellular Senescence. Cancer Res. 73, 3615–3624.

Yaffe, M.P., Noggle, S.A., and Solomon, S.L. (2016). Raising the standards of stem cell line quality. Nat. Cell Biol. 18, 236–237.

Yang, Z., Nakagawa, K., Sarkar, A., Maruyama, J., Iwasa, H., Bao, Y., Ishigami-Yuasa, M., Ito, S., Kagechika, H., Hata, S., et al. (2014). Screening with a Novel Cell-Based Assay for TAZ Activators Identifies a Compound That Enhances Myogenesis in C2C12 Cells and Facilitates Muscle Repair in a Muscle Injury Model. Mol. Cell. Biol. 34, 1607–1621.

Zhao, B., Ye, X., Yu, J., Li, L., Li, W., Li, S., Yu, J., Lin, J.D., Wang, C.-Y., and Chinnaiyan, A.M. (2008). TEAD mediates YAP-dependent gene induction and growth control. Genes Dev. 22, 1962–1971.

Zhao, H., Traganos, F., Dobrucki, J., Wlodkowic, D., and Darzynkiewicz, Z. (2009). Induction of DNA damage response by the supravital probes of nucleic acids. Cytom. Part J. Int. Soc. Anal. Cytol. 75, 510–519.

Zunder, E.R., Lujan, E., Goltsev, Y., Wernig, M., and Nolan, G.P. (2015). A continuous molecular roadmap to iPSC reprogramming through progression analysis of single-cell mass cytometry. Cell Stem Cell 16, 323–337.

